# CD163 and Tim-4 identify resident intestinal macrophages across sub-tissular regions that are spatially regulated by TGF-β

**DOI:** 10.1101/2023.08.21.553672

**Authors:** Ian E. Prise, Vignesh Jayaraman, Verena Kästele, Rufus H. Daw, Kelly Wemyss, Hayley Bridgeman, Sabrina Tamburrano, Patrick Strangward, Christine Chew, Liesbet Martens, Charlotte L. Scott, Martin Guilliams, Antony D. Adamson, Joanne E. Konkel, Tovah N. Shaw, John R. Grainger

## Abstract

In bodily organs, macrophages are localised in poorly understood tissular and sub-tissular niches associated with defined macrophage ontogeny and activity. In the intestine, a paradigm is emerging that long-lived macrophages are dominantly present in the muscular layer, while highly monocyte-replenished populations are found in the lamina propria beneath the epithelial barrier. Whether longevity is restricted in such a simplified manner has not been well explored. Moreover, the impact of specific gut-associated factors on long-lived macrophage functionality and niche occupancy is unknown. We generated sc-RNA-Seq data from wild-type and *Ccr2*^−/−^ mice to identify phenotypic features of long-lived macrophage populations in distinct intestinal niches and identified CD163 as a useful marker to distinguish submucosal/muscularis (S/M) from lamina propria (LP) macrophages. Challenging the emerging paradigm, long-lived macrophages, identified by Tim-4 expression, were found in the LP and S/M. Long-lived LP macrophages are restrained in their response to proinflammatory stimulation compared to short-lived populations in the same location, and to the long-lived population within the S/M. Employing a novel *Timd4*^cre^*Tgfbr2*^fl/fl^ mouse line we demonstrate distinct functions of TGF-β on long-lived macrophages in these two compartments. Importantly, in *Timd4*^cre^*Tgfbr2*^fl/fl^ mice, zonation of CD163^+^ macrophages in the S/M was lost, suggesting TGF-β plays an unappreciated role in positioning of macrophages in the tissue. These data highlight the importance of considering ontogeny and niche when assessing the action of key intestinal regulatory signals.

## Introduction

Molecular and functional heterogeneity of resident tissue macrophages (RTMs) is a topic of longstanding interest due to the range of roles they perform and their potential to be manipulated for therapeutic advantage (*1–4*). Transcriptional and functional heterogeneity of RTMs has been described at the level of ontogeny (*5–8*), tissue site (*9, 10*), and increasingly sub-tissular niche (*11–15*). The implications of this extraordinary level of heterogeneity are in the process of being revealed, but to date include the findings that RTMs fulfil functions tailored to supporting the tissue they reside in (*16–18*) and the cells they are next to (*19, 20*). In turn, the cells creating the spatial niche around the RTM are thought to maintain and instruct the RTM in its function (*21–23*). While several studies have begun to identify relationships between RTM location and function in the lung (*13, 14, 24*), liver (*25*) and brain (*12*), the small intestine remains a site where these relationships are still relatively poorly understood. This is surprising given the diverse environments intestinal RTMs can be found in. For example, the intestine can be broadly divided into four layers: mucosa (including the lamina propria), submucosa, muscularis propria and serosa (*26–29*). In each of these environments, macrophages can be found next to sub-tissular structures, such as nerves (*22, 28, 30*) and blood vessels (*15, 30,31*).

As well as identifying macrophages in physically distinct compartments, intestinal macrophages with different levels of dependency on monocyte replacement have also recently been identified (*30, 32, 33*). In particular, we identified Tim-4 as a marker of intestinal macrophages with low levels of monocyte replenishment. Prior to this, it was thought that all intestinal macrophages were continuously and rapidly replenished from monocytes, largely due to the tonic inflammation in the intestine driven by the high commensal microbial burden (*34, 35*). Whilst there is emerging agreement on the presence of a self-maintaining intestinal macrophage population, there is not yet consensus on the best markers to identify them or the location of these cells. Indeed, the emerging paradigm is that the lamina propria (LP) macrophages present in the tonically inflamed environment beneath the intestinal epithelium are constantly replenished from monocytes, while long-lived macrophages are present in the deeper tissues, such as the submucosa and muscularis (S/M) (*36, 37*). Resulting from the lack of consensus on markers to identify these long-lived macrophages and knowledge about where they are located, the functions and factors regulating long-lived macrophages in distinct sub-tissular regions of the intestine have not been explored.

Long-lived macrophages are of particular interest as similar populations in other organs have been shown to perform functions integral to tissue homeostasis (*16, 18*) and may present a mechanistic link between temporally separated events such as early life disruption to microbial colonisation and the onset of childhood allergy (*38, 39*). Understanding of the location, development and function of long-lived intestinal macrophage subsets is still developing but may provide novel mechanistic targets to maintain and restore intestinal health.

The majority of functional studies on intestinal macrophages have used whole mouse intestine and the conclusions of these studies are often ascribed to LP macrophages, as the more abundant subset (*40, 41*). Increasingly, however, unique transcriptional and functional roles for S/M macrophages are being reported. Muscularis macrophages have been characterised as possessing a more ‘M2’-like phenotype and have been shown to be involved in bidirectional interactions with enteric nerves, controlling peristalsis and protecting tissue and nerves during infection (*20, 22, 28, 42*). To date, many of these studies have relied on the physical splitting of the muscularis layer away from the LP in combination with immunohistochemical imaging using pan-macrophage markers and architectural features of intestinal tissue layers to determine the location and function of muscularis macrophages. Further in-depth functional analyses would benefit from the identification of specific surface markers distinguishing between LP and S/M macrophages, allowing for their functional assessment in mixed populations and for genetic manipulation using subset-specific Cre-recombinase mice.

Here, making use of RNA-Seq and lineage-tracing transgenic animals (*Cx3cr1*^creER^*:R26-yfp*), we report the presence of long-lived Tim-4^+^ macrophages within the LP of the small intestine, as well as the S/M, and establish CD163 as a marker of S/M macrophages. The previously unappreciated CD163^−^Tim4^+^ macrophages located in the LP are restrained in their response to proinflammatory stimulation compared to short-lived populations in the same location. CD163^+^Tim4^+^ macrophages are located in the S/M and produce inducible nitric oxide synthase (iNOS) and high levels of tumour necrosis factor (TNF) in response to proinflammatory factors. Based on this new understanding of macrophage ontogeny in the LP and S/M we generated *Timd4*^cre^ mice to allow specific targeting of these two long-lived macrophage populations. Employing these new mice, we modulated TGF-β-signalling (*Timd4*^cre^*Tgfbr2*^fl/fl^) as a candidate pathway differentially controlled between the two Tim-4^+^ RTM subsets and revealed that TGF-β regulates spatial localisation of CD163^+^ macrophages and iNOS-production in LP RTMs. Taken together our findings go against the current dogma that macrophages in the LP are constantly highly replenished from monocytes and for the first time begin to reveal the functions of key immunoregulatory cytokines on RTMs in distinct intestinal locations.

## Results

### sc-RNA-Seq of wild-type and *Ccr2*^−/−^ animals identifies Tim-4^+^CD163^−^ LP macrophages and Tim-4^+^CD163^+^ S/M macrophages

In order to identify distinct populations of resident tissue macrophages, we began by utilising single-cell (sc)-RNA-Seq of live CD45^+^Lin^−^CD11b^+^MHCII^+^CD64^+^ cells (**Fig. S1A**) from the small intestine of healthy wild-type (WT) and *Ccr2*^−/−^ mice (*43*). *Ccr2*^−/−^ mice have a paucity of recruited circulating monocytes (*44*), and we have previously shown that they are enriched for intestinal long-lived macrophages (*32*). Comparison of the intestinal macrophage populations in these two strains was, therefore, used to help understand the relationship between macrophage populations and their ontogeny.

Following automated clustering combined with manual assessment of gene expression profiles, we initially clustered 5,639 cells into 21 clusters that were interpretable as biologically meaningful (**Fig. S1B**). All the clusters were assessed for expression of the intestinal macrophage markers *Adgre1* (F4/80) and *Cx3cr1* (CX3CR1) (**Fig. S1C**). Given our intention to understand macrophage subsets and ontogeny in health, we excluded clusters that didn’t express high levels of *Adgre1*, and *Cx3cr1* from further analysis (thus clusters 16-20, which were also the smallest, each comprising fewer than 50 cells, were excluded (**Fig. S1C**)). We further excluded clusters that were likely defined by activation/functional states within macrophage subsets rather than subsets in their own right. These were: cluster 11 – proliferating macrophages (*Mki67*); cluster 14 – IFN-activated macrophages (*Ifit1*); and cluster 15 – macrophages that had associated with epithelial cells through doublet formation/phagocytosis (*Spink1*) (**Fig. S1D**). Finally, we excluded clusters 3, 6 and 10 as these cells were absent from WT guts and only present in *Ccr2*^−/−^ guts (**Fig. S1B**), suggesting that they only developed in response to disruption to appropriate macrophage differentiation. Thus, the final sc-RNA-Seq map was generated, containing 10 clusters (**Fig. 1A**).

**Figure 1.**
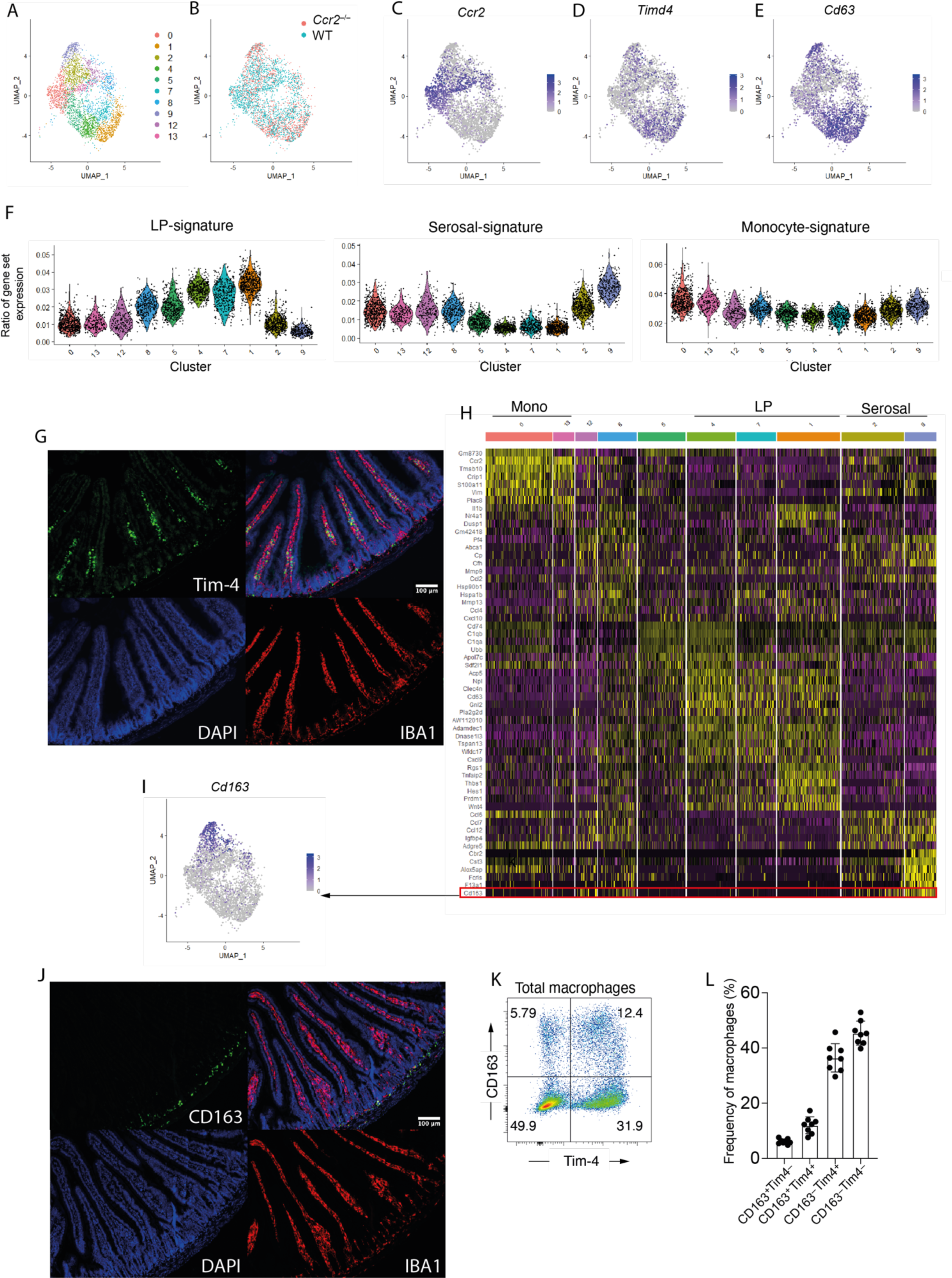
Tim-4 and CD163 expression identify small intestinal macrophages in different sub-tissular locations. **(A)** Uniform manifold approximation and projection (UMAP) plot of sc-RNA-seq data from 3,864 live CD45^+^Lin^−^CD11b^+^MHCII^+^CD64^+^ cells from the small intestine of wild type (WT) C57BL/6 and *Ccr2*^−/−^ mice. Colours denote individual cells assigned to the same cluster. **(B)** A UMAP plot showing the composition of clusters as derived from WT (blue) or *Ccr2*^−/−^ (red) mice. **(C)** A UMAP plot showing expression level of *Ccr2*. **(D)** A UMAP plot showing expression level of *Timd4*. **(E)** A UMAP plot showing expression level of *Cd63.* **(F)** Violin plots showing the similarity score of each cluster against ImmGen datasets for left, lamina propria (LP) macrophages; middle, serosal macrophages; right, monocytes. **(G)** Representative immunofluorescence image of small intestine section from WT mice. DAPI (blue), IBA1 (red), Tim-4 (green). **(H)** Heatmap showing the top differentially expressed genes for all small intestinal macrophage populations identified in Figure 1a, with *Cd163* highlighted. **(I)** A UMAP plot showing expression level of *Cd163.* **(J)** Representative immunofluorescence image of small intestine section from WT mice. DAPI (blue), IBA1 (red), CD163 (green). **(K)** Expression of CD163 and Tim-4 on small intestinal live CD45^+^Lin^−^CD11b^+^MHCII^+^CD64^+^macrophages assessed by flow cytometry. Numbers denote the percentages of cells within the gate. **(L)** Frequency of macrophages expressing Tim-4 and/or CD163 in the small intestine. Error bars show mean ± SD. Data (n = 8) are pooled from eight independent experiments.

We began by analysing the sc-RNA-Seq data to predict the ontogeny of the macrophage clusters. To this end we compared clusters of different macrophage subsets between WT and *Ccr2*^−/−^ mice. Cluster 0 was most distinct from other populations in being almost completely comprised of WT macrophages suggesting that this population is highly dependent on the constant recruitment of monocytes for its generation (**Fig. 1B**). *Ccr2* was enriched in this cluster, and in clusters 8, 12 and 13 (**Fig. 1C**), implying that they also had a strong dependency on monocytes. Contrasting this, clusters 1, 2, 4 and 9 were similarly represented in WT and *Ccr2*^−/−^ animals (**Fig. 1B**), suggesting that these macrophage populations are the least dependent on monocyte recruitment and were, therefore, more long-lived. Supporting this idea, these clusters were enriched for *Timd4* expression (**Fig. 1D**), a marker shown by us, and others, to identify macrophages with lower dependency on monocyte replenishment (*32, 45*). Similarly, *Cd63*, also identified as a marker of intestinal macrophages with low dependency on monocyte replenishment (*30*), was enriched in clusters 1, 2, 4 and 9 (**Fig. 1E**).

In addition to identifying clusters based on their recent development from monocytes, we sought to identify their likely sub-tissular location by comparing their gene signatures with those of the signature of ‘lamina propria (LP)’ and ‘serosal’ macrophages on the ImmGen database (*46*), with ‘serosal’ macrophages likely to also include muscularis macrophages. Clusters 4,7 and 1 scored most highly for ‘lamina propria macrophage’ associated genes and clusters 2 and 9 for ‘serosal macrophage’ associated genes (**Fig. 1F**). Clusters 8 and 5 had an intermediate lamina propria score, suggesting they may be in the processes of transitioning in to mature LP macrophages. In line with their relatively recent entry into the tissue, clusters 0, 13 and 12, enriched for *Ccr2,* were not enriched for either the LP or serosal macrophage gene signature (**Fig. 1F**). As expected, and in line with our previous results (*32*), expression of monocyte-associated genes by any of the clusters was weak, though *Ccr2* enriched clusters, 0 and 13, scored more highly than the rest (**Fig. 1F**). Interestingly, of the clusters which were present in WT and *Ccr2^−/−^* mice, and enriched for expression long-lived self-maintaining RTM genes (*30, 32*), *Timd4* (**Fig. 1D**) and *Cd63* (**Fig. 1E**), clusters 2 and 9 were found to score highly for the ‘serosal macrophage’ signature and 1 and 4 for the ‘LP’ macrophage signal (**Fig. 1F**), suggesting that long-lived macrophages could be found in both sub-tissular locations. These results would, therefore, challenge the current established understanding that long-lived intestinal RTMs are located only within the submucosa/muscularis (*30, 36, 37, 47, 48*).

To validate our sequencing results, we assessed the spatial location of RTMs in the small intestine using immunofluorescence, staining for Tim-4 as a marker of long-lived macrophages (*32*), and found that, in agreement with our sequencing data, Tim-4^+^ macrophages were located in the villi of the LP, with a second, albeit smaller population, also located in the S/M (**Fig. 1G**). In order to identify markers that would distinguish between Tim-4^+^ macrophages in these different locations, we assessed the top differentially expressed genes for each cluster and identified *Cd163* as a highly specific candidate marker of S/M macrophages, enriched in clusters 2 and 9 (**Fig. 1H, I**). Immunofluorescence imaging, as predicted by our sequencing, showed that CD163 expressing macrophages were entirely restricted in their location to the S/M of the small intestine (**Fig. 1J**).

Furthermore, assessment of small intestinal macrophages for CD163 and Tim-4 protein expression showed that they could be separated into four populations: CD163^−^Tim-4^−^, CD163^+^Tim-4^−^, CD163^+^Tim-4^+^, CD163^−^Tim-4^−^ (**Fig. 1K**). The largest macrophage population was the CD163^−^Tim-4^−^ subset (45.4 ± 4.3%), and the smallest population was the CD163^+^ Tim-4^−^ subset (6.2 ± 0.9%) (**Fig. 1L**). These data corroborate, at protein level, that two populations of long-lived Tim-4^+^ macrophages can be identified (CD163^−^ and CD163^+^), likely present in the LP and S/M respectively.

Taken together, these results reveal that Tim-4 may be a marker of longevity, independent of location, while differential expression of CD163 represents a marker reflective of sub-tissular location.

### CD4, Tim-4 and CD163 identify macrophage populations with distinct maturation and tissular signatures

Previously, we and others have suggested that CD4 can be used as a maturation marker for gut macrophages (*32, 49*). In agreement with this *Cd4* was largely absent in cluster 0, the cluster dominated by recently arrived WT *Ccr2*^+^ macrophages, yet present in the majority of other clusters (**Fig. S1E**). We investigated whether this maturation marker could be used across both the CD163^+^ compartment and CD163^−^ compartment to understand ontogeny and maturation. Indeed, assessment of Tim-4, CD163 and CD4 expression allowed identification of 6 subsets (**Fig. 2A**), putatively reflecting longevity, location, and maturation respectively. Notably, CD4^−^ cells were a higher frequency in the LP suggesting there is a greater proportion of high turnover cells in this location (**Fig. 2A**). Reciprocally Tim-4^+^ cells were most frequent within the CD163^+^ compartment (**Fig. 2A**).

**Figure 2.**
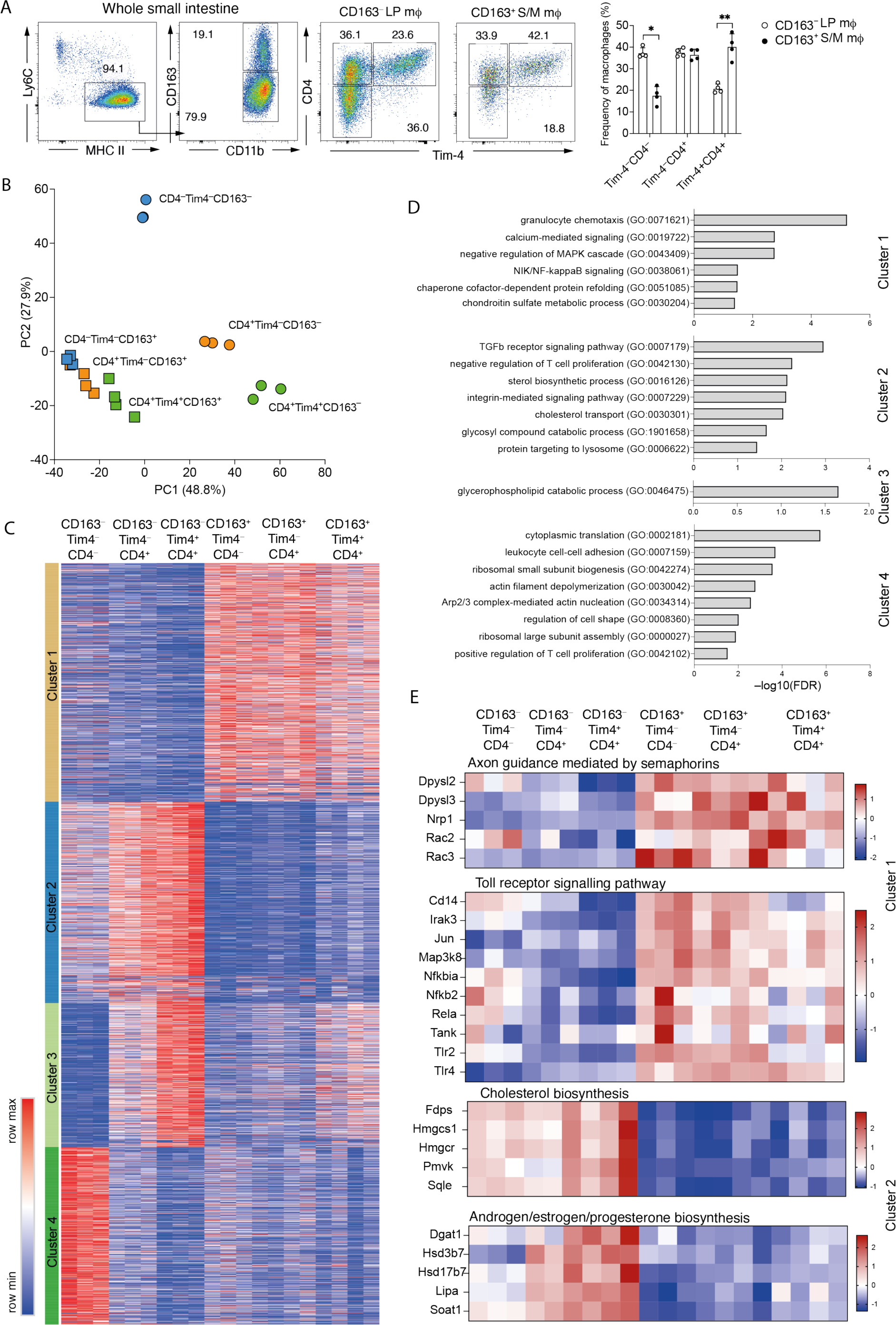
CD4, Tim-4 and CD163 identify transcriptionally distinct macrophages in the small intestine. **(A)** Left: representative flow cytometry plots showing the frequency of Tim-4^−^CD4^−^, Tim-4^−^ CD4^+^, and Tim-4^+^CD4^+^ expressing cells within the CD163^−^ and CD163^+^ subsets of total small intestinal macrophages from WT mice. Right: frequency of Tim-4^−^CD4^−^, Tim-4^−^CD4^+^, and Tim-4^+^CD4^+^ expressing cells within the CD163^−^ and CD163^+^ subsets of total small intestinal macrophages from WT mice. Data (n = 4) are pooled from two independent experiments. **(B)** Principal component analysis (PCA) of global gene expression from CD163^−^ and CD163^+^ subsets of Tim-4^−^CD4^−^, Tim-4^−^CD4^+^, and Tim-4^+^CD4^+^ macrophage of the small intestine isolated by FACS from 5 pooled WT mice, from at least 3 independent sorts. **(C)** Gene expression profile of the 3206 genes differentially expressed (p-adjusted < 0.001) in CD163^−^ and CD163^+^ subsets of Tim-4^−^CD4^−^, Tim-4^−^CD4^+^, and Tim-4^+^CD4^+^ macrophage of the small intestine with clusters identified by k-means clustering. **(D)** Top gene ontology (GO) terms associated with each of the 4 clusters formed by the 3206 differentially expressed genes. **(E)** Heatmaps showing expression profiles of genes in the top two pathways identified by PANTHER Pathway analysis for clusters 1 and 2.

To confirm that identifying cells in this manner could be broadly useful in identifying cells of different maturity and sub-tissular niche we sorted each of these populations and undertook bulk RNA-Seq. Principle component analysis (PCA) revealed that CD163 expression, along with CD163 associated genes, was responsible for the largest separation (PC1) between these 6 macrophage subsets, suggesting that sub-tissular location was the biggest factor controlling differential gene expression of total intestinal macrophages (**Fig. 2B**). The second largest factor (PC2) driving separation of these subsets was the maturity of the cells, as determined by the expression of CD4, though this was only apparent in the CD163^−^ subsets (**Fig. 2B**), suggesting that the acquisition of a ‘lamina propria’ signature takes longer than a ‘submucosal/muscularis’ signature. This analysis confirms that greater heterogeneity is seen within the LP macrophages (CD163^−^) than the S/M macrophages (CD163^+^) (*30*).

Unsupervised hierarchical k-means clustering of 3206 differentially expressed genes (DEGs) (p-adjusted < 0.001) generated 4 clusters (**Fig. 2C and Table S1**). Genes upregulated in all CD163^+^ populations were found in cluster 1 and include *Bmp2, F13a1, Mrc1, Lyve1, Cd36, and Folr2* supporting their identity as submucosal/muscularis macrophages (*14, 22, 28, 50*). This demonstrated that although CD163^+^ macrophages can be split into three populations based on CD4 and Tim-4, all of these populations have a strong muscularis signature (**Fig. 2C and Table S1**).

Similarly, Cluster 2 defined an LP signature evident in all CD163^−^ populations and included *Itgax* (CD11c), a marker previously used to distinguish between lamina propria and muscularis macrophages (*22*). Unlike in the CD163^+^ S/M macrophages Tim-4^−^CD4^−^ cells had lower expression of LP genes suggesting that they were less differentiated (**Fig. 2C**). This concept was re-enforced by Cluster 3 that consisted of genes that were associated with intestinal macrophage maturation and self-maintenance such as *Cd4, Dtx3, Hes1, Timd4 and Cd63* (*30, 32, 49, 51*). The CD4^−^ LP macrophages expressed much lower levels of these genes than CD4^+^ LP macrophages and there was an expression gradient suggesting gradual upregulation of these genes as the cells mature (**Fig. 2C and Table S1**).

Finally, cluster 4 contained genes most highly expressed in CD163^−^CD4^−^Tim4^−^ cells, associated with recent arrival into the intestine and early differentiation from monocyte to macrophage (*Ly6c2, Ccr2, Id3, Bach1, Nfkb1*) (*32, 45, 49, 51*). Again, supporting the idea that the CD4^−^Tim-4^−^CD163^−^ cells are the most recently monocyte derived cells (**Fig. 2C and Table S1**).

Top gene ontology (GO) terms for genes within the CD163^+^ associated cluster 1 were generally related to intracellular signalling (NIK/NF-kappaB signalling, negative regulation of MAPK cascade, calcium-mediated signalling), and leukocyte migration (granulocyte chemotaxis). Top GO terms for cluster 2, associated with CD163^−^ macrophages, were related to protein transport (protein targeting to lysosome) and lipid metabolism (sterol biosynthetic process, cholesterol transport) T cell proliferation (negative regulation of T cell proliferation), TGF-β signalling (transforming growth factor beta receptor signalling pathway), and integrin signalling (integrin-mediated signalling pathway). Only one GO term was associated with cluster 3, suggesting involvement in lipid metabolism. In line with recent monocyte entry into the tissue, cluster 4 was associated with GO terms relating to actin assembly and cell migration (**Fig. 2D**). PANTHER pathway analysis further suggested that CD163^+^ macrophages (cluster 1) are involved in neuron development (axon guidance mediated by semaphorins) and innate immune cell recognition of pathogen associated molecular patterns (Toll receptor signalling pathway), and CD163^−^ (cluster 2) macrophages are involved in lipid synthesis (cholesterol biosynthesis and androgen/estrogen/progesterone biosynthesis) (**Fig. 2E**). No pathways were identified from the genes in cluster 3 and pathway analysis of genes in cluster 4 supported the identification of CD163^−^Tim4^−^CD4^−^ cells as migrating cells (cytoskeleton regulation by Rho GTPase) with inflammatory potential (inflammation mediated by chemokine and signalling pathway) (**Fig. S2**).

Thus, Tim-4, CD4 and CD163 can be usefully employed in tandem to distinguish populations of macrophages of different sub-tissular localisation and also likely ontogeny. However, there are differences in the types of populations these markers will define between the LP and muscularis. In the LP, 3 distinct populations will be defined: 1) CD4^−^Tim-4^−^ that appear to be recently monocyte-derived and poorly differentiated; 2) CD4^+^Tim-4^−^ macrophages that have a strong LP-signature but weaker expression of longevity markers than Tim-4^+^ macrophages; and 3) CD4^+^Tim-4^+^ macrophages that express the highest levels of longevity markers. Contrasting this, in the muscularis although CD163^+^CD4^+^ and CD4^−^ are evident by flow-cytometry these have almost indistinguishable transcriptional profiles and both express strong muscularis gene signatures. However, they are potentially shorter-lived than Tim-4^+^CD163^+^ macrophages as they express reduced levels of genes more associated with longevity e.g. *Cd63*.

### Long-lived Tim-4^+^ macrophages are present in the LP and S/M

As discussed, an emerging paradigm in the field is that LP macrophages are short-lived and replenished from monocytes while macrophages in the S/M are locally maintained (*36, 37*). Our sc-RNA-Seq data and bulk RNA-Seq data implied that differential macrophages of various maturation states and longevity were present in both LP and S/M and that these could be distinguished on a bulk population level by CD163, CD4 and Tim-4.

Based on this we employed CD163, CD4 and Tim-4 to assess longevity in both the LP and S/M. To this end adult *Cx3cr1*^creER^*:R26-yfp* mice were dosed with tamoxifen for 5 consecutive days, as previously described (*32, 52*). At 5 days, 16, 24 and 32 weeks after the final tamoxifen dose, mice were sacrificed to assess the expression of YFP within CD163^+^ and CD163^−^ macrophage subsets.

At 5 days after cessation of tamoxifen treatment, YFP expression within CD163^+^ macrophages (Tim-4^−^CD4^−^, Tim-4^−^CD4^+^ and Tim-4^+^CD4^+^) was uniformly >70%, on average, and similar to that seen in two of the CD163^−^ populations (Tim-4^−^CD4^+^ and Tim-4^+^CD4^+^) (**Fig. 3A**). Labelling of CD163^−^Tim-4^−^CD4^−^ macrophages was, however, lower than seen in the all-other subsets, likely due to their more recent differentiation from monocytes, associated with more recent upregulation of CX3CR1 (*53*). By 32 weeks, the majority of Tim-4^−^ macrophages, both CD163^+^ and CD163^−^ had been replaced by YFP^−^ cells (**Fig. 3B**). As suggested by our sequencing data, Tim-4^+^ macrophages still comprised a significant proportion of YFP^+^ cells, regardless of CD163 expression status, and are, therefore, likely present in the LP and S/M (**Fig. 3B**). This confirms the utility of Tim-4 as a marker of self-maintaining macrophages in different compartments of the small intestine, and at longer time points than we previously showed (*32*).

**Figure 3.**
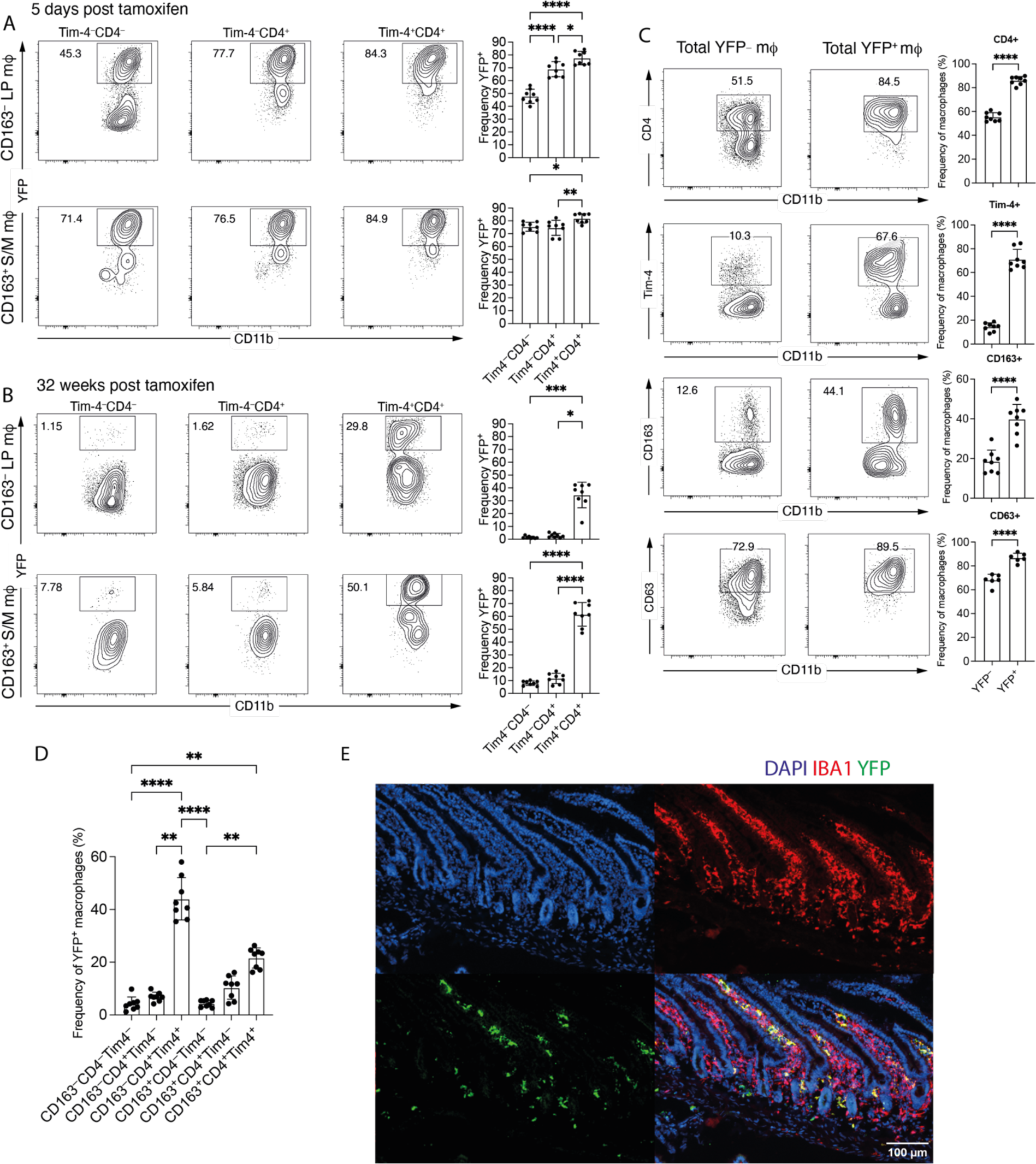
Long-lived macrophages are present in the LP and S/M. **(A)** Left: representative flow cytometry plots showing the frequency of YFP-expressing cells within the CD163^−^ and CD163^+^ subsets of Tim-4^−^CD4^−^, Tim-4^−^CD4^+^, and Tim-4^+^CD4^+^ macrophage of the small intestine of *Cx3cr1*^creER^*:R26-yfp* mice, 5 days after tamoxifen treatment. Right: frequency of YFP-expressing cells within the CD163^−^ and CD163^+^ subsets of Tim-4^−^CD4^−^, Tim-4^−^CD4^+^, and Tim-4^+^CD4^+^ macrophage of the small intestine of *Cx3cr1*^creER^*:R26-yfp* mice, 5 days after tamoxifen treatment. **(B)** Left: representative flow cytometry plots showing the frequency of YFP-expressing cells within the CD163^−^ and CD163^+^ subsets of Tim-4^−^CD4^−^, Tim-4^−^CD4^+^, and Tim-4^+^CD4^+^ macrophage of the small intestine of *Cx3cr1*^creER^*:R26-yfp* mice, 32 weeks after tamoxifen treatment. Right: frequency of YFP-expressing cells within the CD163^−^ and CD163^+^ subsets of Tim-4^−^CD4^−^, Tim-4^−^CD4^+^, and Tim-4^+^CD4^+^ macrophage of the small intestine of *Cx3cr1*^creER^*:R26-yfp* mice, 32 weeks after tamoxifen treatment. **(C)** Left: representative flow cytometry plots showing expression of CD4, Tim-4, CD163, or CD63 in YFP^+^ and YFP^−^ macrophages of the small intestine from *Cx3cr1*^creER^*:R26-yfp*, 32 weeks after tamoxifen treatment. Right: frequency of YFP^+^ and YFP^−^ macrophages expressing CD4, Tim-4, CD163, or CD63 in the small intestine of *Cx3cr1*^creER^*:R26-yfp* mice, 32 weeks after tamoxifen treatment. **(D)** Frequency of total YFP^+^ macrophages expressing CD163, CD4 and Tim-4 in the small intestine of *Cx3cr1*^creER^*:R26-yfp* mice, 32 weeks after tamoxifen treatment. **(E)** Representative immunofluorescence of small intestine section from *Cx3cr1*^creER^*:R26-yfp*, 32 weeks after tamoxifen treatment. DAPI (blue), IBA1 (red), YFP (green). **(A – C)** Numbers in flow cytometry plots denote the percentages of cells within the gate. **(A – D)** Data (n = 6 - 8) are pooled from 2-3 independent experiments. Error bars show mean ± SD. Statistical comparisons between 2 groups were performed with an unpaired t test with Welch’s correction for parametric data and a Mann Whitney test for nonparametric data. Statistical comparisons between more than 2 groups were performed with a with one-way ANOVA, with Tukey’s multiple comparison test for parametric data and a Kruskal Wallis test with Dunn’s multiple comparison test for nonparametric data. *, P ≤0.05; **, P ≤0.01; ***, P ≤0.001; and ****, P ≤0.0001.

At 16, 24, and 32 weeks, CD163 expression further differentiated the Tim-4^+^ macrophages, with CD163^+^Tim4^+^ macrophages retaining a higher frequency of YFP^+^ cells than their CD163^−^ counterparts (**Fig. 3B and Fig. S3A**). CD163 expression also differentiated the Tim-4^−^ macrophage subsets at these time points, though YFP expression was not more than 24%, on average, in any of these populations (**Fig. S3A**). These findings were further confirmed by the generation of shielded bone marrow chimeras, as previously described (*32, 54*) (**Fig. S3B**).

When gating on total YFP^+^ macrophages at 32 weeks, we observed that they were characterised by higher expression of CD4, Tim-4 and CD163 than total YFP^−^ macrophages (**Fig. 3C**), suggesting that they are useful markers of longevity. CD63, a marker recently identified as a potential marker of long-lived, self-renewing intestinal RTMs (*30*) was most highly expressed by YFP^+^ macrophages but was also expressed by YFP^−^ macrophages (**Fig. 3C**), potentially limiting its utility as a specific marker of long-lived intestinal RTMs.

Furthermore, when total YFP^+^ macrophages at 32 weeks were assessed for expression of CD163, CD4 and Tim-4, the greatest proportion of YFP^+^ macrophages were found to be CD163^−^CD4^+^Tim-4^+^ (44.0 ± 8.0%), followed by CD163^+^CD4^+^Tim-4^+^ (21.6 ± 3.8%) (**Fig. 3D**). This led us to predict that a striking proportion of long-lived macrophages would be located within the LP, as well as the S/M, and not just the S/M as previously suggested (*30*). We, therefore, examined the location of YFP^+^ macrophages within the small intestine of mice, 32 weeks after tamoxifen treatment and found that these macrophages were indeed located in both sub-tissular compartments (**Fig. 3E**).

These data demonstrate that in both the LP (CD163^−^) and the submucosa/muscularis (CD163^+^), Tim-4 identifies the macrophages with the lowest dependency on monocytic replenishment. Although the longest-lived macrophages were those Tim-4^+^ cells present in the submucosa/muscularis, even at 32 weeks there was a substantial frequency of Tim-4^+^ macrophages in the LP that had not turned over from monocytes, demonstrating that there are long-lived macrophages in this region of the tissue.

### Small intestinal macrophages differ in responsiveness to stimulation, according to their phenotype and location

Much of our understanding of the function of intestinal RTMs has been based on whole mixed preparations of digested lamina propria and muscularis (*40, 41*), prior to our understanding of long-lived intestinal macrophages (*30, 32*). Alternatively, muscularis macrophages have been isolated utilising a gut splitting approach (*20, 28*).

Making use of our new marker of location (CD163) and longevity marker (Tim-4) we sought to establish whether there were differences between the functions of the longest-lived macrophage populations in the LP and the muscularis in a mixed culture. To this end single cell suspensions from the intestine were cultured with pro-inflammatory stimuli and assessed for expression of pro-IL-1β, TNF-α and iNOS.

We initially focussed on whether location would impact responsiveness to proinflammatory stimuli. For long-lived (Tim-4^+^) macrophages, location did not significantly impact production of pro-IL-1β in the presence of LPS (**Fig. 4A**). Short-lived LP macrophages (Tim-4^−^), however, were more likely to produce pro-IL-1β than their S/M counterparts (**Fig. 4A**). Contrasting this, S/M macrophages expressed higher levels of both TNF-α and iNOS than LP macrophages, for both long and short-lived populations (**Fig. 4B, C**). Thus, the sub-tissular location from which intestinal RTMs come from has an impact on their propensity to produce proinflammatory factors, when equally exposed to stimuli in a mixed culture system.

**Figure 4.**
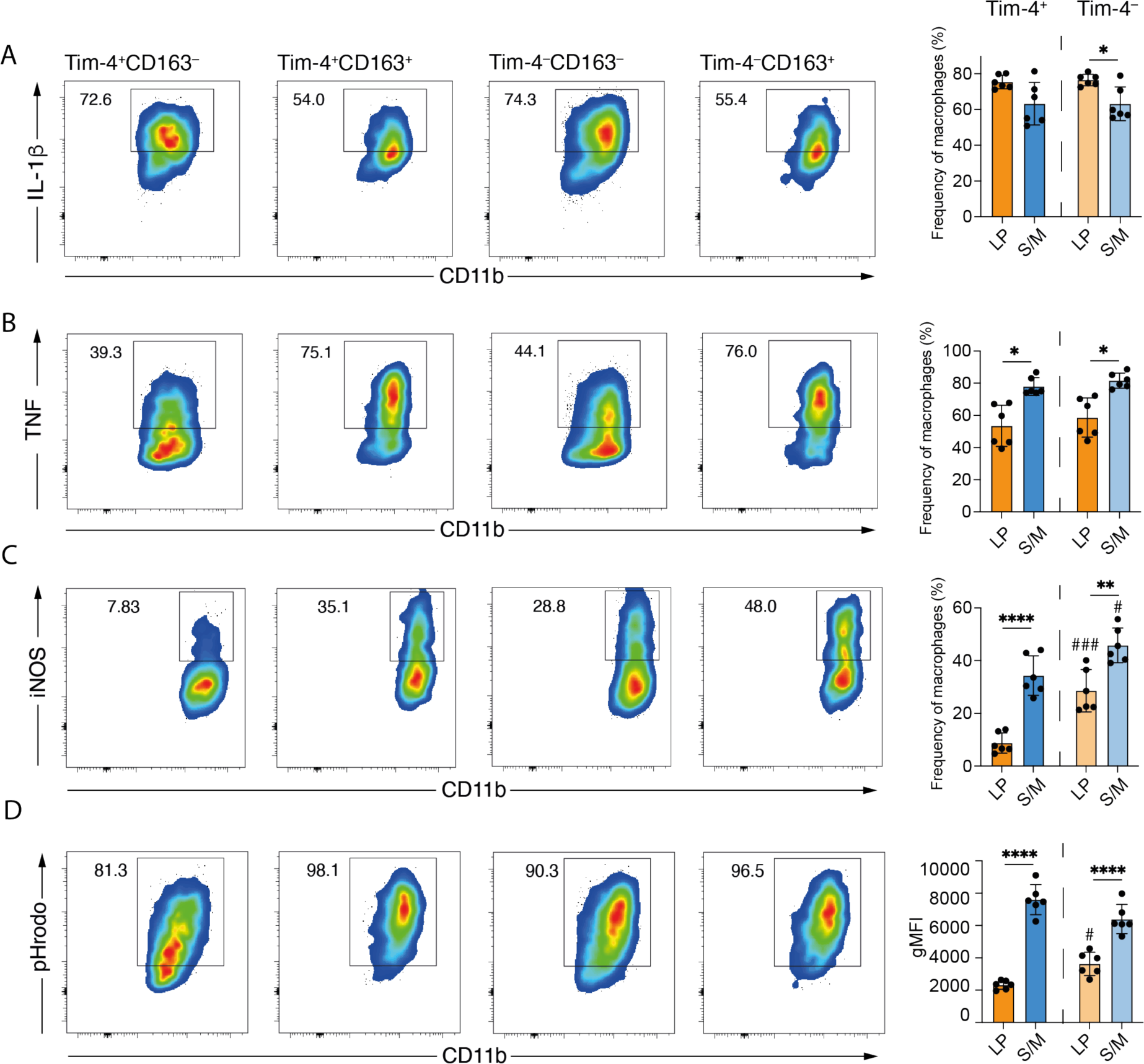
Sub-tissular location is a dominant determinant of small intestinal macrophage function. **(A)** Left: representative flow cytometry plots showing the frequency of cells expressing pro-IL-1β from small intestinal cells enriched for macrophages and stimulated for 3 hours. Right: frequencies of macrophages expressing pro-IL1β. **(B)** Left: representative flow cytometry plots showing the frequency of cells expressing TNF from small intestinal cells enriched for macrophages and stimulated for 3 hours. Right: frequencies of macrophages expressing TNF. **(C)** Left: representative flow cytometry plots showing the frequency of cells expressing iNOS from small intestinal cells enriched for macrophages and stimulated overnight. Right: frequencies of macrophages expressing iNOS. **(D)** Left: representative flow cytometry plots showing the frequency of pHrodo^+^ cells after 40 minutes incubation. Right: geometric mean fluorescence intensity (gMFI) of macrophages for pHrodo expression. **(A - D)** Numbers in flow cytometry plots denote the percentages of cells within the gate. Error bars show mean ± SD. Data (n = 6) are pooled from at least 2 independent experiments. Statistical comparisons were performed with a with one-way ANOVA, with Tukey’s multiple comparison test for parametric data and a Kruskal Wallis test with Dunn’s multiple comparison test for nonparametric data. Significance is shown for comparisons between Tim-4^+^ and Tim-4^−^ macrophages within the same compartments *, P ≤0.05; **, P ≤0.01; and ****, P ≤0.0001, and between Tim-4^+^ and Tim-4^−^ macrophages across compartments #, P ≤ 0.01; ###, P ≤ 0.001.

As well as effects of sub-tissular location, we were also able to investigate functionality of shorter-lived macrophage populations (Tim-4^−^), compared with long-lived populations (Tim-4^+^) in the same cultures. Production of TNF and pro-IL-1β was independent of longevity (**Fig. 4A, B**). However, iNOS production was enhanced in Tim-4^−^ populations in both the LP and muscularis, suggesting that longevity or time spent in the tissue, as well as sub-tissular location, is an important determinant of iNOS production (**Fig. 4C**). For iNOS, this resulted in long-lived LP macrophages producing strikingly low levels of iNOS, compared with the other 3 macrophage subsets (**Fig. 4C**).

Phagocytic capacity is another important functional feature of intestinal macrophages (*33, 34*). We directly assessed phagocytosis *ex vivo* by culturing single cell suspensions with pHrodo bioparticles. As for proinflammatory cytokine production, detectable differences in phagocytic capacity were evident between Tim-4^+^ LP (CD163^−^) and S/M (CD163^+^) macrophages, with LP macrophages displaying a reduced phagocytic capacity (**Fig. 4D**). In comparison to location, longevity had a smaller effect on phagocytic capacity, and only within the LP compartment, highlighting location as the major factor determining differences in phagocytic functionality (**Fig. 4D**).

Thus, when activated under the same conditions, detectable differences in macrophage function can be observed based on both location and longevity; iNOS in particular is impacted by both location and longevity in the LP.

### Distinct sub-tissular requirements for TGF-β-signalling in long-lived intestinal macrophages

In the intestine, a number of signals have been identified that are crucial to instruction of intestinal macrophage differentiation including TGF-β, IL-10, lipid mediators and short-chain fatty acids (*49, 55, 56*). However, the impact of the same intestinal specific signals on different resident macrophages in sub-tissular compartments has not been explored; in part because it was not appreciated that long-lived macrophages are present in both the LP and S/M. Our bulk RNA-Seq data (**Fig. 2**) allowed us to identify TGF-β receptor signalling pathways as a GO term (GO-0007179) that was differentially enriched between LP and S/M (**Fig. 2D**). We further investigated the genes that contribute to GO-0007179, identifying known negative and positive regulators. Interestingly, although enriched in the LP (**Fig. 2C, D**), a range of genes involved in TGF-β receptor signalling were found to be differentially expressed in CD163^−^ and CD163^+^ subsets, including those involved in negative (*Smad6*, *Smad7*) and positive (*Furin, Thbs1*) regulation of this signalling pathway (**Fig. 5A**). These results imply that RTMs in both the LP and S/M receive TGF-β signals, and that they exhibit distinct TGF-β transcriptional responses, likely due to differences in regulation of the TGF-β signal transduction system within their respective sub-tissular locations. We, therefore, chose to further investigate TGF-β signalling as a pathway that could have important effects across the LP and S/M compartments.

**Figure 5.**
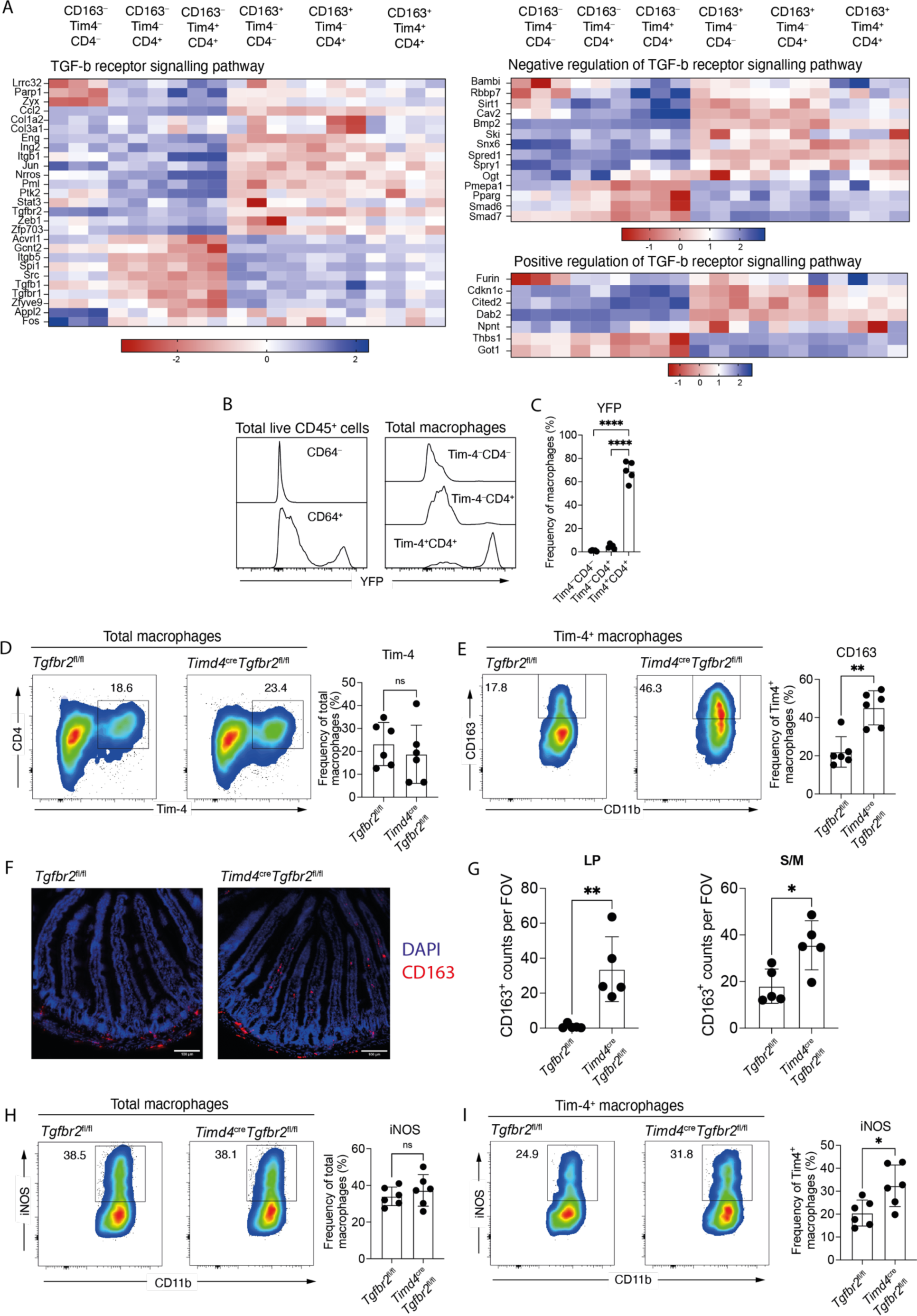
TGF-β signalling controls the localisation and responsiveness of long-lived small intestinal macrophages. **(A)** Heatmaps showing expression profiles of genes listed under the TGF-β receptor signalling pathway GO term (GO-0007179) and identified as a DEG by bulk RNA-seq. **(B)** Left: expression of YFP in CD64^−^ and CD64^+^ cells from total live CD45^+^ cells in the small intestine. Right: expression of YFP in Tim-4^−^CD4^−^, Tim-4^−^CD4^+^ and Tim-4^+^CD4^+^ macrophages in the small intestine. **(C)** Frequency of small intestinal Tim-4^−^CD4^−^, Tim-4^−^CD4^+^ and Tim-4^+^CD4^+^ macrophages expressing YFP. **(D)** Left: representative flow cytometry plots showing the frequency of small intestinal macrophages expressing Tim-4 in *Timd4*^cre^*Tgfbr2*^fl/fl^ and *Tgfbr2^fl/^*^fl^ mice. Right: frequencies of Tim-4^+^ macrophages in the small intestine of *Timd4*^cre^*Tgfbr2*^fl/fl^ and *Tgfbr2^fl/^*^fl^ mice. **(E)** Left: representative flow cytometry plots showing the frequency of Tim-4^+^ macrophages expressing CD163. Right: frequencies of Tim-4^+^ macrophages expressing CD163. **(F)** Representative immunofluorescence images of the small intestine showing the location of CD163 expressing cells in *Timd4*^cre^*Tgfbr2*^fl/fl^ and *Tgfbr2^fl/^*^fl^ mice. DAPI (blue), CD163 (red). **(G)** Quantification of CD163 expressing cells in the LP (left) and S/M (right) of the small intestine in *Timd4*^cre^*Tgfbr2*^fl/fl^ and *Tgfbr2^fl/^*^fl^ mice, by field of view (FOV). Data (n = 5 per group) are pooled from 2 independent experiments. **(H)** Left: representative flow cytometry plots showing the frequency of total macrophages expressing iNOS in response to overnight LPS and IFNγ stimulation. Right: frequencies of total macrophages expressing iNOS in response to overnight LPS and IFNγ stimulation. **(I)** Left: representative flow cytometry plots showing the frequency of Tim-4^+^ macrophages expressing iNOS in response to overnight LPS and IFNγ stimulation. Right: frequencies of Tim-4^+^ macrophages expressing iNOS in response to overnight LPS and IFNγ stimulation **(D, E, H, I)** Numbers in flow cytometry plots denote the percentages of cells within the gate. Data (n = 6 per group) are pooled from 2 independent experiments. **(C – E, G – I)** Error bars show mean ± SD. **(B, C, G)** Data (n = 5) are pooled from 2 independent experiments. Statistical comparisons were performed with an unpaired t test with Welch’s correction for parametric data and a Mann Whitney test for nonparametric data. *, P ≤0.05; **, P ≤0.01; ****, P ≤0.0001.

Currently, no transgenic model exists to easily target long-lived macrophages across sub-tissular compartments in the gut. Using available approaches, such as *Cx3cr1*^cre^ mice it is not possible to selectively target Tim-4^+^ long-lived intestinal macrophages. We, therefore, generated a new *Timd4*^cre^ mouse. Although this mouse will impact long-lived macrophages at many sites (*45*), in the gut it will allow for selective targeting of the *Timd4*^+^ cells while sparing other populations. To confirm the specificity of the *Timd4*^cre^, we crossed this transgenic animal to the *R26-yfp* mouse (*57*), in which all cells that have expressed *Timd4*^cre^ will be labelled with YFP. As expected YFP was highly expressed by intestinal CD64^+^ macrophages, and not CD64^−^ cells, and was well expressed by Tim-4^+^ macrophages (**Fig 5B, C**). Assessment of blood and bone marrow monocytes revealed no YFP expression, and high levels of YFP expression were seen in liver Tim-4^+^ Kupffer cells and Tim-4^+^ large peritoneal macrophages,but not Tim-4^−^ large peritoneal macrophages or small peritoneal macrophages (**Fig. S4A**), thus confirming the specificity of our novel *Timd4*^cre^ mouse.

Given our interest in TGF-β signalling in the LP and S/M*, Timd4*^cre^ animals were crossed with *Tgfbr2^fl/fl^* mice (*58*) to delete TGF-β-receptor 2 (TGF-βR2) on long-lived Tim-4^+^ macrophages in both compartments. Following the loss of TGF-βR2 on Tim-4^+^ macrophages, we observed no overt signs of spontaneous inflammation in the small intestine (**Fig. S4B**) or colon (**Fig. S4C**), and no change in frequency of small intestinal Tim4^+^ macrophages (**Fig. 5D**). Within the Tim-4^+^ population, however, the frequency of CD163^+^ macrophages was increased in *Timd4*^cre^*Tgfbr2*^fl/fl^ mice compared with *Tgfbr2^fl/fl^* controls (**Fig. 5E**).

The increase of CD163-expressing Tim-4^+^ macrophages in the small intestine of *Timd4*^cre^*Tgfbr2*^fl/fl^ animals suggested that in the absence of TGF-β-signalling, long-lived submucosa/muscularis macrophages were selectively expanded. To explore this, we imaged CD163-expressing macrophages in the small intestine of *Timd4*^cre^*Tgfbr2*^fl/fl^, compared to their *Tgfbr2*^fl/fl^ littermates (**Fig. 5F**). Unexpectedly, the expansion of CD163^+^ cells was associated only with a modest increase in numbers of CD163^+^ cells in the muscularis with most of the expansion being driven by the loss of zonation (restriction) of CD163^+^ macrophages and their *de novo* presence in the LP (**Fig. 5G**). Thus, in the small intestine TGF-β-signalling on long-lived Tim-4^+^ macrophages is critical to ensure normal distribution of CD163^+^ macrophages.

In **Fig. 4C**, we showed that long-lived LP macrophages are less likely to produce iNOS than short-lived LP macrophages, or S/M macrophages. As long-lived LP macrophages in *Timd4*^cre^*Tgfbr2*^fl/fl^ mice upregulate CD163 expression, usually associated with S/M macrophages, we asked whether long-lived LP macrophages also adopted functional similarities to S/M macrophages, such as increased iNOS production.

On global macrophages there was no difference in iNOS production between *Timd4*^cre^*Tgfbr2*^fl/fl^ mice and *Tgfbr2^fl/fl^* controls (**Fig. 5H**). When we assessed only Tim-4^+^ long-lived macrophages, however, we observed an increase in iNOS production in *Timd4*^cre^*Tgfbr2*^fl/fl^ (**Fig. 5I**), with the iNOS being produced mostly by the CD163^+^ macrophages (**Fig. S4D**). As expected, as this was a *Timd4*^cre^ mouse, no increase in CD163 expression or iNOS induction was seen in Tim-4^−^ macrophages (**Fig. S4E**).

Thus, using our new phenotyping and transgenic approaches to explore the distinct tissular niches of long-lived gut macrophages, our data reveal an unappreciated role for TGF-β in defining zonation of CD163^+^ macrophages in the S/M and a role in regulating iNOS production in the LP layer.

## Discussion

We are only just beginning to understand the diversity of RTM ontogeny and functionality in tissue and sub-tissular compartments. Here we have demonstrated Tim-4^+^ macrophages with distinct low-replenishment from monocytes in the LP and S/M. Until relatively recently it was thought that all macrophages in the gut were constantly replenished from monocytes (*34*). In our previous publication (*32*), we did not establish where Tim-4^+^ macrophages were located within the intestinal tissue and based on subsequent publications it was assumed that these cells were largely in the S/M (*30*). Indeed, a perspective has even been raised that monocyte replenishment rates in the LP are so high this should not be considered as a niche (*36*). Notably, publications from Kelsall *et al*. (*50*) and Chiaranunt *et al*. (*59*) suggested that Tim-4^+^ macrophages may also be in multiple sub-tissular niches within the colon, but this was not fully explored with respect to LP versus S/M localisation.

Making use of sc-RNA-Seq from WT mice and *Ccr2*^−/−^ mice, which have a paucity of monocyte-derived macrophages, we identified candidate populations of Tim-4^+^ long-lived macrophages in both the LP (CD163^−^) and S/M (CD163^+^). Employing markers of longevity (Tim-4), maturity (CD4) and tissue localisation (CD163) in *Cx3cr1*^creER^*:R26-yfp* mice we found that, up to 32 weeks post tamoxifen treatment, macrophages expressing YFP were still present in both the LP and S/M. While the S/M macrophages were the longest-lived, a substantial proportion of LP macrophages were still expressing YFP at 32 weeks. Thus, in contrast to the prevailing paradigm, macrophages in the tonically inflamed environment of the LP can persist for long periods of time. This finding is in contrast with a recent paper showing long-lived RTMs only in the S/M (*30*). We cannot currently explain this difference, as the mouse model and method used to label and detect long-lived RTMs were the same. As more groups become aware of, and study, these long-lived intestinal RTMs, a consensus on their location should emerge. Infectious and inflammatory insults are known to cause depletion of macrophages in other organs, with repopulation occurring by engraftment of newly recruited macrophages or proliferation of remaining macrophages (*23*). After such an event, it is possible that long-lived macrophages that would otherwise have been present in the LP would be replaced predominantly by newly recruited macrophages. This still does not negate the fact that clearly there are contexts where LP macrophages can exist in their tissue environment for long periods of time; it is now crucial to investigate what environmental factors support longevity of LP macrophages and the importance of their longevity to intestinal mucosal health.

In addition to demonstrating the presence of long-lived Tim-4^+^ macrophages in the LP, we also identified CD163 as an unappreciated marker, in the small intestine, of macrophages restricted to the S/M (**Fig. 1**). CD163 is a scavenger receptor that acts by endocytosing haemoglobin-haptoglobin complexes (*60*). Its expression has been associated with perivascular macrophages in mice, including those in the brain (*61*) as well as tumour associated macrophages (TAMs) (*62*). Whether S/M CD163^+^ macrophages in the gut are localised with blood vessels is not yet understood but the identification of CD163 as a marker to distinguish between LP and S/M macrophages will provide new opportunities for selective targeting. For example, employing *Cd163*^cre^ animals (*62*) to target S/M macrophages, with the caveat that vascular macrophages in other organs will be impacted.

Mapping of our findings of long-lived LP and S/M macrophages onto human macrophage populations remains to be undertaken. While Tim-4 itself is unlikely to be a conserved marker for human long-lived gut macrophages, based on its lack of expression in recent datasets (*63–65*), the overall signatures observed may be more useful. Similarly, CD163 is reported to be expressed by macrophages within the villi of the human intestine (*66*), limiting the utility of this marker to identify regionally restricted RTMs to murine models. Aligning our data to human single cell data sets may allow for potential alternative and conserved markers to be identified, such as the recently identified FOLR2, which was found to be expressed in submucosal and muscularis macrophages in human colonic samples (*65*), and also identified to be a conserved marker of self-sustaining macrophage populations across tissues and species (*45*). In line with these reports, *Folr2* was identified as an upregulated DEG in cluster 1 (CD163^+^ macrophages) in our bulk RNA-seq data (**Table S1**).

Irrespective of the precise conserved factors between mice and humans, identification of long-lived Tim-4^+^ macrophages in both the LP and S/M provided us with an opportunity to understand how the gut-associated signal TGF-β acts across the two compartments. It is well established that global loss of TGF-β1 signalling causes multiorgan inflammation, including in the stomach and colon (*67, 68*), and intestinal RTMs have been shown to require TGF-β signalling for their organ specific differentiation (*49*). However, studies looking at the effect of disrupting TGF-β signalling, specifically in myeloid cells, on the intestine have produced conflicting results (*69–74*), and none have assessed how TGF-β signalling influences intestinal RTMs in different sub-tissular compartments or RTMs with different longevities. Due to the lack of tools available to target intestinal RTMs and, in particular, specific subsets, we generated a *Timd4*^cre^ mouse, which allowed us to specifically target Tim-4^+^ macrophages.

Using this new transgenic mouse, we identified differential effects of TGF-β signalling on two similarly long-lived Tim-4^+^ macrophage subsets, within different sub-tissular regions of the small intestine. TGF-β signalling was found to regulate CD163 expression and iNOS production by long-lived Tim-4^+^ macrophages in the LP, with loss of TGF-β signalling in these cells resulting in aberrant expression of these phenotypic and functional markers in long-lived LP macrophages. The consequences of the presence of dysregulated long-lived macrophages in the LP remain to be explored but a predisposition to inflammatory disease is a possibility and may provide a mechanistic link between disrupted TGF-β signalling and IBD (*75*).

Differences in regulation of long-lived RTMs in the LP and S/M raise the possibility that TGF-β, which is produced and stored as an inactive complex, is more abundant within the LP compared with the S/M, or that it is activated or utilised differently by long-lived Tim-4^+^ RTMs, within these compartments. Indeed, the magnitude and duration of TGF-β signalling are tightly controlled at multiple levels, including synthesis, release from its inactive form, receptor expression and stability of downstream signalling molecules, including negative regulatory elements (*76*). Any number, or combination, of these tightly regulated mechanisms may be responsible for the observed differences in response to TGF-β signalling between long-lived Tim-4^+^ macrophages in the LP and S/M. Early studies looking at the regional distribution of TGF-β protein within the murine intestine suggest that epithelial cells are a major source of TGF-β1, particularly at the villus tips of the small intestine and surface epithelium of the colon (*77, 78*). Barnard *et al.*, additionally described moderate TGF-β1 expression within the muscularis mucosa of the small intestine and colon (*77*). Studies in humans have reported greater *TGF-β* mRNA expression in the LP than epithelial cells (*79, 80*), though expression in the muscularis mucosa was not examined. In contrast to the early murine studies, and more similar to the human reports, a more recent study in mice shows higher *Tgf-β1* mRNA expression in the LP than in the epithelium, or the muscularis mucosa, during homeostasis (*81*). This paper identified macrophages themselves as an important source of TGF-β1 and our bulk RNA-sequencing data similarly suggests that CD163^−^ LP RTMs express higher levels of *Tgfb1* than CD163^+^ S/M RTMs (**Fig. 5A and Table S1**).

When taken together, it is tempting to speculate, that our data suggest two very distinct pathways for macrophage development in the small intestine, depending on sub-tissular location. Movement of monocytes into the LP is associated with a sequential acquisition of an LP macrophage-signature associated with length of time present in the tissue and relating to CD4 and Tim-4 expression (**Fig. 2B**). In the longest-lived macrophages, ongoing exposure to TGF-β leads to a hyporesponsive phenotype following exposure to IFN-γ/LPS (**Fig. 4C**). Contrasting this, macrophages in the S/M more rapidly adopt a transcriptional profile that does not further change with time spent in the tissues, as determined by upregulation of CD4 and Tim-4 (**Fig. 2B**), and is associated with maintained responsiveness to IFN-γ/LPS stimulation (**Fig. 4C**). Establishing the environmental factors determining which pathway a monocyte takes upon entry into the intestine will require further investigation. Overall, our new understanding of distinct long-lived macrophages will open-up new routes for researching intestinal macrophage populations and inform understanding of signalling pathways that could be targeted for the treatment of intestinal inflammatory diseases.

## Materials and Methods

### Mice

Male C57BL/6J (CD45.2) were purchased from Charles River Laboratories, strain #:000664, (experiments within the UK) or Janvier (experiment within Belgium) and housed in individually ventilated cages under specific pathogen-free conditions. The following mice, originally from The Jackson Laboratory, were bred in-house under the same conditions; *Ccr2^−/−^*, strain #:004999 (*43*), *Cx3cr1*^creER^, strain #:020940 (*53*), *R26R-yfp*, strain #:006148 (*57*), *TbRII*, JAX stock #:012603 (*58*), congenic CD45.1, strain #:002014 (*82*). *C57BL/6J.Timd4^Em1Uman^* mice were generated as described below, by Antony D. Adamson, at the University of Manchester. All experiments were approved by The University of Manchester or Edinburgh Local Ethical Review Committee and were performed in accordance with the UK Home Office Animals (Scientific Procedures) Act 1986.

### Generation of C57BL/6J.Timd4^Em1Uman^ mice

To express Cre recombinase under the control of *Timd4* regulatory regions, but preserve Tim-4 expression, we used CRISPR-Cas9 to integrate Cre recombinase immediately downstream of the *Timd4* translation stat site, along with a T2A self-cleaving peptide, to ensure both genes/proteins are generated from the locus.

The sgRNA sequences, which target the Timd4 ATG (GATCCTATCAAAATGTCCAA and AGCCCCTTGGACATTTTGAT) were purchased as full length Alt-R sgRNA oligos (Integrated DNA Technologies) and resuspended in sterile, RNase free injection buffer (TrisHCl 1mM, pH 7.5, EDTA 0.1mM). For our donor repair template we used the EASI-CRISPR long-ssDNA strategy (*83*) and generated a homology flanked lssDNA donor using protocols described in Bennett *et al*. 2021 (*84*).

For embryo microinjection the annealed sgRNA was complexed with Cas9 protein (New England Biolabs) at room temperature for 10 minutes, before addition of long ssDNA donor (final concentrations; sgRNA 20 ng/μl, Cas9 protein 20 ng/μl, lssDNA 10 ng/μl). CRISPR reagents were directly microinjected into the nuclei of C57BL/6J (Envigo) zygotes using standard protocols. Zygotes were cultured overnight, and the resulting 2 cell embryos surgically implanted into the oviduct of day 0.5 post-coitum pseudopregnant mice.

Potential founder mice were screened by PCR, first using primers that flank the sgRNA sites (JG01_F gccaccatgagaaaagtgcct, JG01_R tccccaaacacccaaatcca), which both identifies editing activity in the form of InDels from NHEJ repair and can also detect larger products, implying integration of the Cre-T2A through Homology directed repair (HDR). Secondary PCRs used the same primers in combination with Cre specific primers (Cre_F gatcgctgccaggatatacg, Cre_R gtgccttctctacacctgcg). Candidate founders giving positive products in all three PCR reactions were further characterised by amplifying again with the JG01 F/R primers using high fidelity Phusion polymerase (NEB), the larger product gel extracted and subcloned into pCRblunt (Invitrogen) and Sanger sequenced with M13 Forward and Reverse primers. Alignment of the sequencing confirmed integration of the tag. A single founder was bred with WT C57BL/6J and germline transmission confirmed with the same array of assays.

### Tamoxifen treatment

Tamoxifen (Sigma-Aldrich) was dissolved in 10% ethanol and 90% corn oil to a concentration of 50 or 100 mg/ml. Male and female were dosed with 5 mg by oral gavage for five consecutive days.

### Generation of shield chimeras

Male CD45.2^+^ host mice aged 6–8 weeks were anaesthetized by intraperitoneal administration of ketamine (80 mg/kg; Vetoquinol) and xylazine (8 mg/kg; Bayer). Anaesthetized mice were positioned beneath a lead sheet shielding the lower two thirds of the body, including the intestine, from a split dose of irradiation (2× 5.5 Gy). Mice therefore received partial body irradiation with only the head, thorax, and forelimbs left exposed. After recovery from anasthesia, mice were reconstituted by intravenous injection with 2 × 10^6^ CD90.2^+^ T cell–depleted donor BM cells from congenic CD45.1^+^ WT donor animals. T cells were depleted using CD90.2 microbeads (Miltenyi Biotec). Mice were maintained on 0.03% enrofloxacin in drinking water for up to 1 wk before and for 2 wk after irradiation and then were housed in autoclaved cages with sterile water, diet, and bedding. Reconstitution was allowed to occur for 24 weeks before analysis.

### Tissue preparation and cell isolation

#### Small intestine preparations

Cells were isolated as previously described with some modifications (Sun et al., 2007). In brief, after dissection of the small intestine and colon, Peyer’s patches were removed from the length of the small intestine, and both small intestine and colon were cut longitudinally and washed thoroughly with PBS on ice. Subsequently, to remove intestinal epithelial cells and leukocytes, small intestines and colons were cut into segments (2–3 cm) and incubated in prewarmed media (RPMI 1640) supplemented with penicillin and streptomycin, 3% FCS, 20 mM HEPES, 100 U/ml polymyxin B (Sigma-Aldrich), 5 mM EDTA, and 1 mM freshly thawed dithiothreitol for 15 min at 37°C with agitation. After incubation, gut segments were repeatedly shaken in fresh serum-free media with 2 mM EDTA and 20 mM HEPES to ensure optimal dissociation of intestinal epithelial cells and leukocytes. Remaining tissue (lamina propria and muscularis) was minced and digested at 37°C for 30 min with continuous stirring in serum-free RPMI containing 20 mM HEPES, 0.1 mg/ml liberase TL (Roche), and 0.5 mg/ml DNase. Digested tissue was passed sequentially through a 70-µm and 40-µm cell strainer, and after pelleting, it was resuspended in RPMI 1640 supplemented with 2mM L-glut, 1× NEAA, 1mM sodium pyruvate, 20mM HEPES, 1× penicillin and streptomycin, and 10% FCS until staining.

#### Blood

Blood was collected into EDTA-coated syringes from sacrificed mice. Suspensions were washed and resuspended in ACK lysing buffer (Gibco) for 3 min on ice, twice. Suspensions were then washed and resuspended in RPMI 1640 supplemented with 2mM L-glut, 1× NEAA, 1mM sodium pyruvate, 20mM HEPES, 1× penicillin and streptomycin, and 10% FCS until staining.

#### Bone marrow

Bone marrow was collected from hind-leg femurs, cut at one end, and placed inverted into collection tubes pulsed in a microfuge. Cells were passed through a 70-µm filter and resuspended in ACK lysing buffer (Gibco) for 3 min on ice. Suspensions were then washed and resuspended in RPMI 1640 supplemented with 2mM L-glut, 1× NEAA, 1mM sodium pyruvate, 20mM HEPES, 1× penicillin and streptomycin, and 10% FCS until staining.

#### Peritoneal exudate cells

Peritoneal cavity exudate was collected by injecting 8 mL PBS into the peritoneal cavity. The resulting cell suspension was then reclaimed, washed, and resuspended in RPMI 1640 supplemented with 2 mM L-glut, 1× NEAA, 1 mM sodium pyruvate, 20 mM HEPES, 1× penicillin and streptomycin, and 10% FCS until staining.

#### Liver

Liver tissue finely minced in pre-warmed digest media (RPMI 1640), supplemented with 20 mM HEPES, 1 mg/mL Collagenase IV (Gibco), and 10 μg/mL DNAse (Sigma-Aldrich) and incubated for 20 minutes at 37°C with continuous stirring. Post-incubation, the digested livers were passed sequentially through a 70-µm and 40-µm cell strainer, and after pelleting, were resuspended in RPMI 1640 supplemented with 2 mM L-glut, 1× NEAA, 1 mM sodium pyruvate, 20 mM HEPES, 1× penicillin and streptomycin, and 10% FCS until staining.

### In vitro assays

For stimulations, whole small intestine, prepared as above, was pelleted and resuspended in 5 mL of nycoprep (Axis Shield) or OptiPrep (STEMCELL Technologies), prepared according to the manufacturer’s directions. 2 mL of FCS-free RPMI was carefully layered on top and the preparation was centrifuged at 800g for 15 minutes at room temperature with the brake set to ‘low’. Mononuclear phagocytes at the interface were collected into a fresh tube, counted and 1 × 10^^6^ cells were seeded for stimulations.

#### Pro-IL-1b and TNF induction

1 × 10^^6^ cells were incubated in RPMI 1640 supplemented with 2mM L-glut, 1× NEAA, 1mM sodium pyruvate, 20mM HEPES, 1× penicillin and streptomycin, and 10% FCS in a 96 well ‘U’ bottomed plate and stimulated with 1μg/mL LPS-EB Ultrapure (Invivogen) and Brefeldin A (eBioScience). After 3 hours, cells were washed and stained with antibodies for assessment by flow cytometry.

#### iNOS induction

1 × 10^^6^ cells were seeded in a non-TC treated 24 well plate, in RPMI 1640 supplemented with 2 mM L-glut, 1× NEAA, 1 mM sodium pyruvate, 20 mM HEPES, 1× penicillin and streptomycin, 50 mg/mL gentamicin, and 10% FCS. Cells were incubated at 37°C and 5% CO2 for one hour to settle and then stimulated with 20 ng/mL recombinant IFNg (BioLegend), 100 ng/mL LPS-EB Ultrapure (Invivogen) and 20 ng/mL recombinant murine M-CSF (PeproTech) overnight. The next day, cells were collected from the supernatant and detached from the plate using ACCUTASE (STEMCELL Technologies) for assessment by flow cytometry.

#### Phagocytosis assay

1 × 10^^6^ cells from whole small intestine, prepared as above, were incubated in RPMI 1640 supplemented with 2 mM L-glut, 1× NEAA, 1 mM sodium pyruvate, 20 mM HEPES, 1× penicillin and streptomycin, and 10% FCS in a 96 well ‘U’ bottomed plate for 30 minutes before the addition of 10 mL of pHrodo Red S. aureus BioParticles Conjugate for Phagocytosis, prepared according to manufactures guidelines (Thermo Fisher Scientific). After 40 minutes, cells were washed and stained with antibodies for assessment by flow cytometry.

### Flow cytometry

Single cell suspensions, prepared as described above, were washed with PBS and stained with the LIVE/DEAD Fixable blue or Near-IR Dead Cell Stain kit (Thermo Fisher Scientific) to exclude dead cells. Subsequently, cells were stained in the dark for 15 min at 4°C with fluorochrome-conjugated antibodies in PBS containing anti-CD16/CD32 (2.4G2; BioXcell or BioLegend). Cells were washed and, in some cases, immediately acquired live, or alternatively, cells were fixed in 2% paraformaldehyde (Sigma-Aldrich) for 10 min at room temperature and resuspended in PBS for later acquisition. Cells were stained with CD4 (RM4-5), CD11b (M1/70), CD11c (N418), CD45 (30F11), CD45.1 (A20), CD45.2 (104), CD64 (X54-5/7.1), CD115 (AFS98), MHCII (I-A/I-E; M5/114.15.2), and Tim-4 (RMT4-54) from BioLegend as well as CD163 (TNKUPJ) and Ly6C (HK1.4) from Thermo Fisher Scientific. The lineage antibody cocktail for excluding lymphocytes and granulocytes included Siglec F (E50-2440) from BD and TCRβ (H57-597), B220 (RA3-6B2), and Ly6G (1A8) from BioLegend. For intracellular antibody staining, fixed cells were permeabilised with Permeabilization Buffer (Thermo Fisher Scientific) and stained with CD63 (NVG-2) and CD206 (C068C2) from BioLegend and pro-IL-1b (NJTEN3), TNF (MP6-XT22) and iNOS (CXNFT) from Thermo Fisher Scientific. Cell acquisition was performed on an LSR Fortessa running FACSDIVA software (BD). For each intestinal sample, typically 10,000–20,000 macrophages were collected. Data were analyzed using FlowJo software (TreeStar).

#### Fluorescence activated cell sorting

Single-cell suspensions for small intestine were prepared as above. Before FACS, on a FACSAria Fusion (BD), isolated cells were suspended in RPMI supplemented with 2% FCS and 2 mM EDTA. Sorted cells were collected in RLT buffer (QIAGEN) supplemented with 2-mercaptoethanol (Sigma) and stored on dry ice before storage at −80°C for subsequent RNA extraction.

#### Bulk RNA sequencing

RNA was extracted from 25,000 – 50,000 cells using an RNeasy micro kit (QIAGEN), following the manufacturer’s instructions. RNA quality was checked using a LabChip GX RNA pico kit (Perkin Elmer) or Bioanalyzer RNA 6000 pico kit (Agilent) and samples were found to have RIN values above 7. RNA sequencing libraries were prepared by Edinburgh Genomics using a SMART-Seq® v4 PLUS RNA Kit (Takara Bio USA) with 9 cycles of amplification. Library cDNA quality was assessed by HSDNA kit on a 2100 Bioanalyzer (Agilent). 16-24 million reads were obtained from each sample using a NovaSeq 6000 platform (Illumina). Reads were filtered with Trimmomatic (v0.36)(*85*). Filtered fastq files were aligned to the mouse GENCODE genome (GRCm38.p5) using STAR (v2.5.3)(*86*). Filtered reads were then sorted, compressed and unaligned reads removed using Samtools (v1.3)(*87, 88*). Aligned reads were then counted, normalized and compared, respectively, using the Cuffquant, Cuffnorm and Cuffdiff function of Cufflinks (v2.2.2)(*89–91*). Heatmaps were visualized using (https://software.broadinstitute.org/morpheus) (*92*). Genes were clustered using one minus Pearson correlation k-means clustering. Gene ontology and pathway analyses were conducted with Panther (http://pantherdb.org). Bulk RNA sequencing data were deposited in the Gene Expression Omnibus public database under accession no. GSE232645.

### Single cell RNA sequencing

#### Single cell isolation and sequencing

Single-cell suspensions of 3 pooled small intestines of male C57BL/6J or 4 pooled small intestines of male *Ccr2*^−/−^ mice were FACS purified for live CD45^+^ Ly6C^−^ Ly6g^−^ TCRb^−^ CD3^−^ B220^−^ Siglec F^−^ CD11b^+^ MHC II^+^ CD64^+^ macrophages. Gene expression libraries were prepared from single cells using the Chromium Controller and Single Cell 3ʹ Reagent Kit v2 (10x Genomics) according to the manufacturer’s protocol to generate single-cell Gel Bead-in-EMulsion (GEMs). Single-cell RNA-Seq libraries were prepared using GemCode Single-Cell 3ʹGel Bead and Library Kit (10x Genomics) according to the manufacturer’s instructions. Indexed sequencing libraries were generated using the reagents in the GemCode Single-Cell 3ʹ Library Kit and sequenced using the Illumina NextSeq500 platform. Sequencing was performed at the VIB Nucleomics Core (Leuven, Belgium) (C57BL/6J) or at the University of Manchester Genomic Technologies Core Facility (Manchester, UK) (*Ccr2*^−/−^). The .bcl sequence data were processed for quality control purposes using bcl2fastq software (v2.20.0.422), and the resulting .fastq files were assessed using FastQC (v0.11.3), FastqScreen (v0.9.2), and FastqStrand (v0.0.5) before pre-processing with the CellRanger pipeline.

### Data processing

Cells with >10% mitochondrial reads were filtered out. The remaining cells were normalized, integrated (using IntegrateData), and analyzed with Seurat (v4.1.1). Geneset module enrichment is presented as a proportion of the transcript count of the genes comprising the geneset module within the total transcript count of each cell and visualized using FeaturePlot (Seurat v4.1.1). sc-RNA sequencing data were deposited in the Gene Expression Omnibus public database under accession no. GSE234018.

#### Immgen database comparison

For comparison of single cell RNA sequencing data to lamina propria and serosal macrophage, and monocyte signatures from the ImmGen database, the Immgen V1 microarray dataset was used (*46*). Genes that were expressed more than 2-fold in MF_103-11b+_SI relative to MF_11cloSer_SI were selected as the Lamina Propria (LP) gene signature and genes that were expressed more than 2-fold in MF_11cloSer_SI relative to MF_103-11b+_SI as the Serosa gene signature. Finally, genes that were expressed more than 2-fold in Mo_6C+II-_Bl relative the mean of MF_103-11b+_SI and MF_11cloSer_SI macrophages were selected as the Classical Monocyte gene signature. Violin plots show the sum of the transcripts of the genes within the gene signatures as a ratio of the total transcripts (transcript per million of gene set of interest/transcript per million of gene set from Immgen datasets).

### Histology and Immunofluorescence microscopy

#### Sample preparation and storage

Small intestines were flushed with PBS, followed by 4% PFA. Short intestinal tissue sections and Swiss-rolls were drop-fixed in 4% PFA for 16-24 hours at 4°C and subsequently embedded and stored in paraffin blocks or drop-fixed in 4% PFA containing 30% sucrose for 3 - 4 hours at 4°C and subsequently snap frozen in OCT in isopentane on dry ice and stored in at –80°C.

#### Histology

Paraffin sections of 5 μM were subjected to normal deparaffination and hydration and stained with haematoxylin solution, Gill no. 3 (Sigma-Aldrich) and eosin Y (Sigma-Aldrich), mounted with DPX (VWR), and cover slipped.

#### Immunofluorescence

Paraffin sections of 8 μM were subjected to normal deparaffination and hydration. Antigen retrieval was carried out using Tris-EDTA pH 9. Tissues were blocked for endogenous biotin using an avidin biotin blocking kit (BioLegend), according to the manufacturer’s instructions and incubated overnight at 4^0^C with one of the following primary antibodies; rat anti-Tim-4 (RMT4-54, BioLegend) or rabbit anti-CD163 (ab182422, abcam), diluted in TBS supplemented with 1% BSA + 0.3% Triton X. The following day, tissues were treated with 0.03 – 0.3% H_2_O_2_ to block endogenous peroxidase before incubation for 1 hour at room temperature with biotin conjugated antibodies against the species of the primary antibody; biotin anti-rat, biotin anti-rabbit. Tissues were subsequently incubated for 1 hour at room temperature with HRP-conjugated streptavidin, followed by Tyramide 555, prepared according to the manufacturer’s instructions (Thermo Fisher Scientific). Tissues stained with CD163 were then subject to antigen retrieval using sodium citrate pH6 and incubated overnight at 4°C with rabbit anti-IBA1 (ab178846, abcam), followed by anti-rabbit AF488 secondary antibody or with rabbit anti-IBA1 AF647 (ab225261, abcam). Tissues stained with Tim-4 were incubated overnight at 4°C with rabbit anti-IBA1 (ab178846, abcam), followed by anti-rabbit AF488 secondary antibody or with rabbit anti-IBA1 AF647 (ab225261, abcam). Finally, tissues were counterstained with DAPI, mounted in prolong diamond mountant (Thermo Fisher Scientific) and coverslipped.

Frozen sections of 10 μM were baked at 50°C for 40 minutes and subsequently incubated with chicken anti-GFP (ab13970, abcam) and rabbit anti-IBA1 (ab178846, abcam) at 4°C overnight. The following day, tissues were blocked for endogenous avidin and biotin using an avidin biotin blocking kit according to the manufacturer’s instructions (BioLegend). Sections were then incubated with biotin conjugated goat anti-chicken (Vector Labs, BA-9010) for an hour at room temperature followed by streptavidin-Cy3 and a goat anti rabbit AF647 for another hour. Finally, tissues were counterstained with DAPI, mounted in prolong diamond mountant (Thermo Fisher Scientific) and coverslipped.

Images were collected on a Zeiss Axioimager.D2 upright microscope using a 10x air objective and captured using a Coolsnap HQ2 camera (Photometrics) through Micromanager software v1.4.23 or on a Nikon Ti2 Eclipse widefield live imaging microscope on a 20x air objective and captured using a Photometrics Prime 95B CMOS camera. Images were processed using ImageJ (*93*).

For quantification of CD163^+^ cells, the area of staining for CD163/µM^2^ was calculated and normalized to the DAPI signal from 5 images per section. Number of cells per mm^2^ were counted using QuPath (*94*). Initially all the images were loaded, and the nuclei were segmented based on DAPI using the ‘Watershed Cell Detection algorithm’ with a threshold of 100 and cell expansion of 3.142 on a full image annotation. An object classifier was then created and trained using 4 random images for detecting CD163 using the RandomTrees Classifier to detect positive cells for CD163. The above steps were batched and applied to all images and exported as a CSV file for further analysis.

### Statistical analyses

Statistical analyses were performed using Prism 9 (GraphPad Software). A Shapiro Wilk test was used to test for normality. For parametric data, two experimental groups were compared using a Student’s *t* test, for paired data, or a Student’s *t* test with Welch’s correction for unpaired data. For nonparametric data, two experimental groups were compared using a Mann Whitney test. Where more than two groups were compared, a one-way ANOVA with Tukey’s multiple comparison test was used for parametric data and a Kruskal Wallis test with Dunn’s multiple comparison test for nonparametric data. Significance was set at P ≤ 0.05.

## Supplementary Materials

**Figs. S1 to S4**

**Table S1**

## Acknowledgements

Thanks go to the following core facilities at The University of Manchester: Bioimaging, Histology, Flow Cytometry, Genome Editing Unit, Genomics Technologies Core Facility, Bioinformatics Core Facility, and Biological Services Facilities. Irradiation in these experiments was performed with assistance from Epistem Ltd. Thanks go to the following core facilities at The University of Edinburgh: Sequencing was carried out by Edinburgh Genomics, which is partly supported with core funding from NERC (UKSBS PR18037); The Centre Optical Instrumentation Laboratory (COIL), which is supported by a Core Grant (203149) to the Wellcome Centre for Cell Biology; the Flow Cytometry and Cell Sorting Facility in the King’s Buildings, the Histology Facility in the Shared University Research Facilities (SuRF), the Institute for Regeneration and Repair Flow Cytometry Facility, and the Bioresearch and Veterinary Services. Thanks go to the VIB Nucleomics, VIB single cell and VIB-UGent Flow Cytometry core facilities.

## Funding

BBSRC grant BB/S01103X/2 (TNS)

Wellcome Trust-University of Edinburgh Institutional Strategic Support Fund 204804/Z/16/Z (TNS)

Senior Fellowship awarded by The Kennedy Trust for Rheumatology Research (JRG)

## Author contributions

Conceptualization: TNS, JRG

Methodology: TNS, IEP, VK, VJ, ST, PS, CLS, ADA

Formal analysis: TNS, IEP, LM

Investigation: TNS, IEP, VK, VJ, KW, HB, ST, RHD, PS, CC, CLS

Resources: MG, ADA, JEK, JRG Data curation: TNS, IEP, LM Project administration: TNS, JRG

Writing – original draft: TNS, JRG

Writing – review & editing TNS, IEP, VK, CLS, JEK, JRG Funding acquisition: TNS, JRG

## Competing interests

Authors declare they have no competing interests.

## Data and materials availability

sc-RNA sequencing data were deposited in the Gene Expression Omnibus public database under accession no. GSE234018.

Bulk RNA sequencing data were deposited in the Gene Expression Omnibus public database under accession no. GSE232645.

All other raw data available on request.

## Notes

https://www.ncbi.nlm.nih.gov/geo/query/acc.cgi?acc=GSE232645

https://www.ncbi.nlm.nih.gov/geo/query/acc.cgi?acc=GSE234018

### Competing Interest Statement

The authors have declared no competing interest.

