## Supplementary Materials for "CD163 and Tim-4 identify resident intestinal macrophages across sub-tissular regions that are spatially regulated by TGF-β"

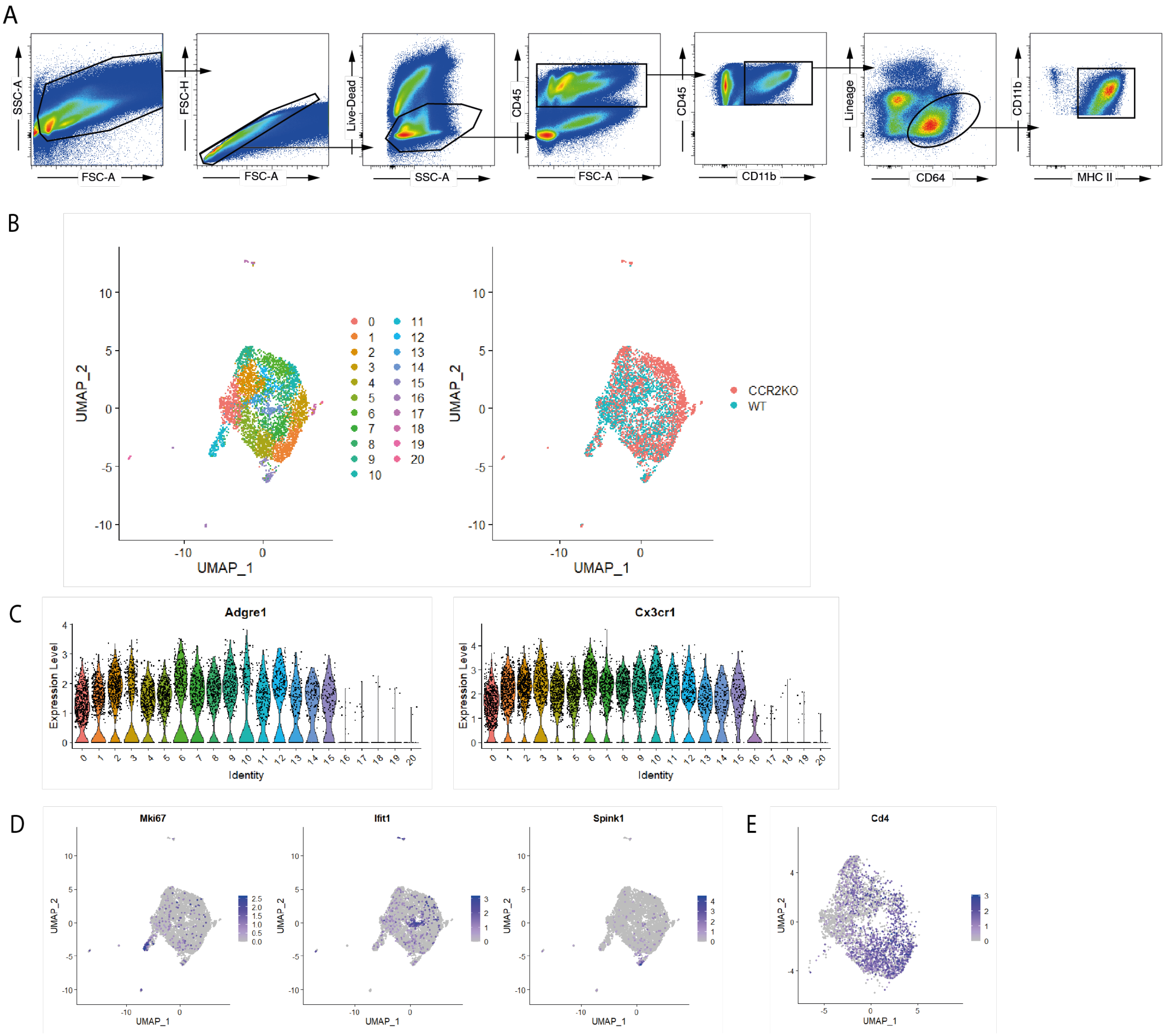


**Supplementary Figure 1. Identification and curation of sc-RNA-seq clusters from WT and Ccr2$^{-/-}$ mice.**

**(A)** Flow cytometry gating strategy for identification of small intestinal macrophages. **(B)** UMAP plots of sc-RNA-seq data from 5,639 live CD45$^{+}$Lin$^{-}$CD11b$^{+}$MHCII$^{+}$CD64$^{+}$ cells from the small intestine of wild type (WT) C57BL/6 and *Ccr2*$^{-/-}$ mice. Left: colours denote individual cells assigned to the same cluster. Right: colours denote cells derived from WT (blue) or *Ccr2*$^{-/-}$ (red) mice. **(C)** Violin plots showing expression of *Adgre1* (F4/80) and *Cx3cr1* (CX3CR1) in each cluster. **(D)** UMAP plots showing expression levels of *Mki67, Ifit1* and *Spink1*. **(e)** UMAP plot showing expression levels of *Cd4*.

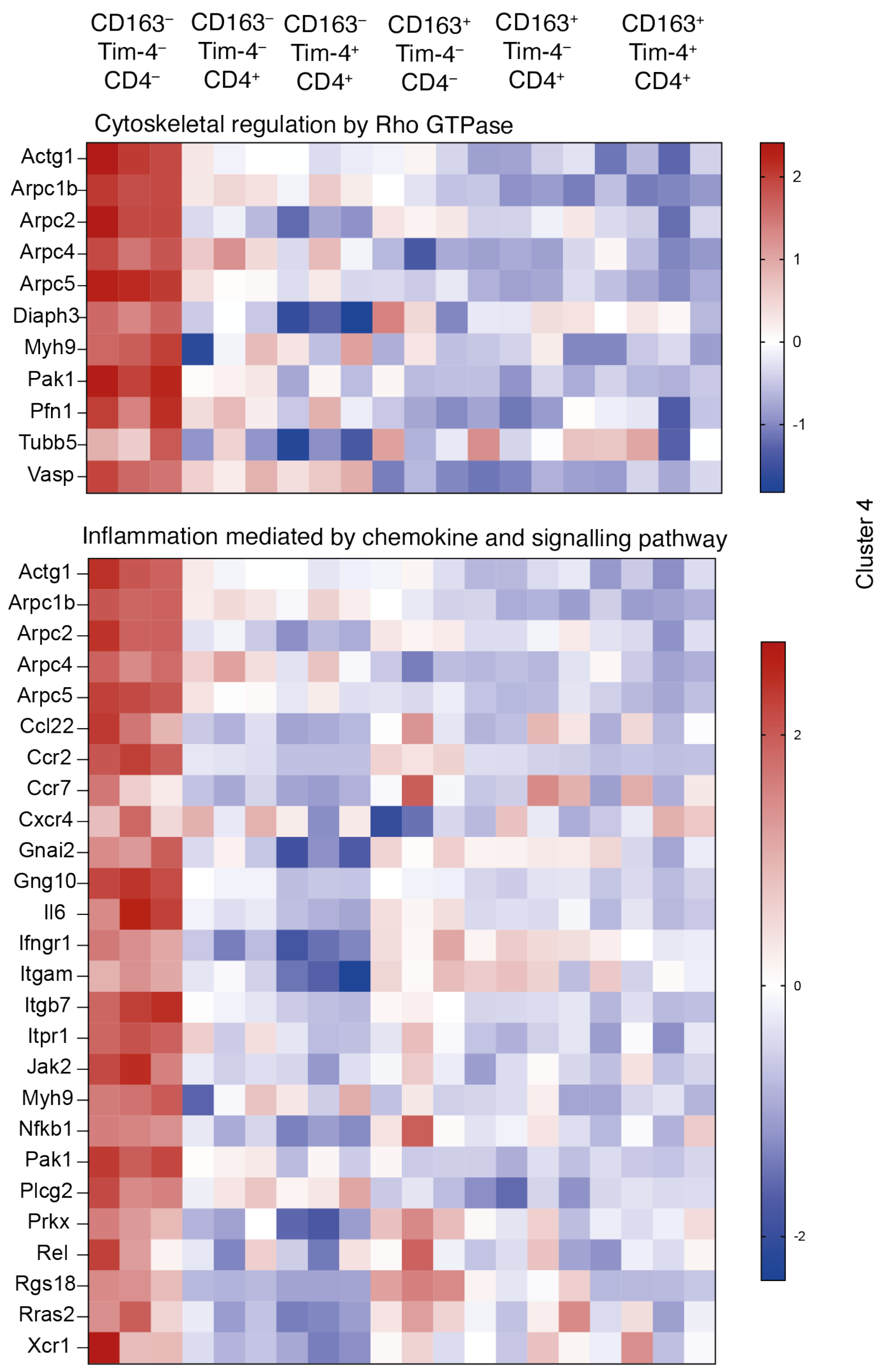


**Supplementary Figure 2. Gene set pathway analysis for cluster 4**

Heatmaps showing expression profiles of genes in the top two pathways identified by PANTHER Pathway analysis for cluster 4. CD163$^{-}$ and CD163$^{+}$ subsets of Tim-4$^{-}$CD4$^{-}$, Tim-4$^{-}$CD4$^{+}$, and Tim-4$^{+}$CD4$^{+}$ macrophage of the small intestine were isolated by FACS from 5 pooled 8 –10-wk-old C57BL/6 WT mice, from at least 3 independent sorts, for bulk RNA-seq.

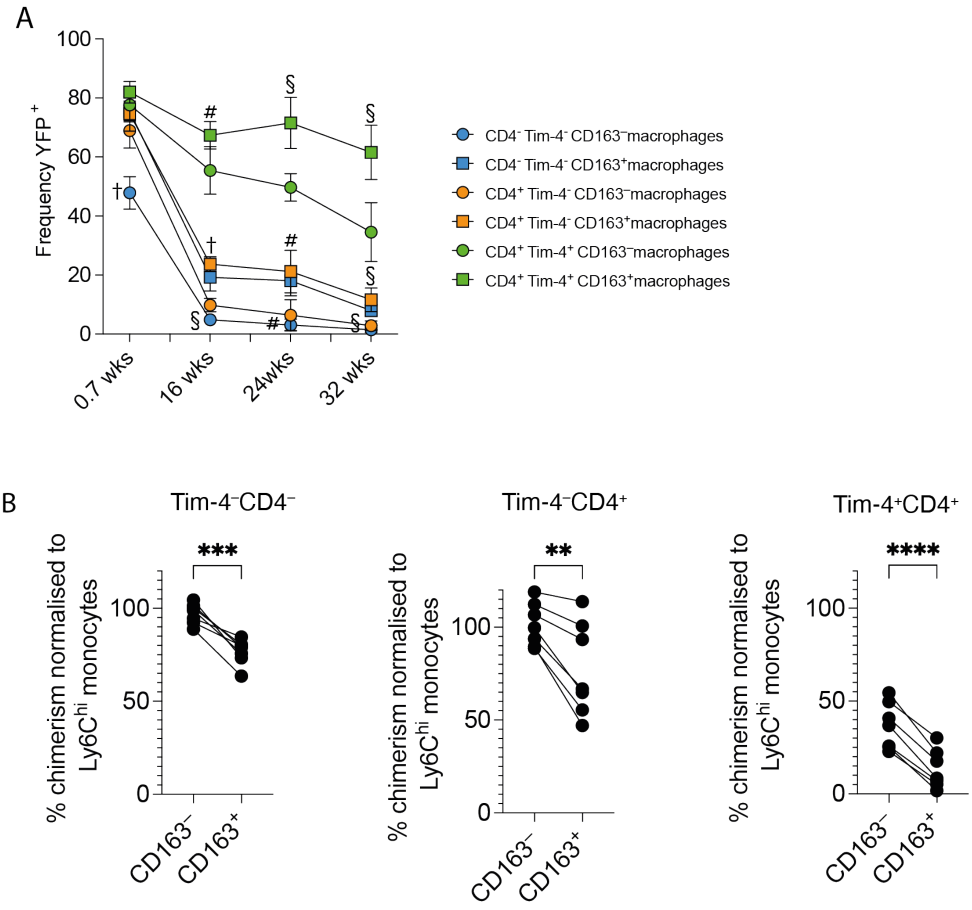


**Supplementary Figure 3. Determination of monocyte contribution to small intestinal macrophage subsets in fate-mapping reporter mice and gut-shielded chimeras.**

**(A)** Frequency of YFP-expressing small intestinal macrophages at 0.7 (5 days), 16, 24 and 32 weeks after tamoxifen treatment of *Cx3cr1*$^{creER}$*:R26-yfp* mice. Data (n = 6 - 8 per group) are pooled from 2 (16 and 24 weeks) or 3 (0.7 and 32 weeks) independent experiments. **(B)** Frequency of donor-derived cells in small intestinal macrophages from abdomen shielded chimeras, 24 weeks after irradiation and reconstitution with congenic bone marrow cells, normalized to the chimerism of Ly6C$^{hi}$ blood monocytes. Data (n = 7 per group) are pooled from 3 harvests, from two independently generated chimera cohorts. **(C)** Error bars show mean ± SD. Statistical comparisons were performed with an unpaired t test with Welch's correction for parametric data and a Mann Whitney test for nonparametric data. Significance is shown for comparisons between CD163$^{+}$ and CD163$^{–}$ counterparts within the Tim-4$^{–}$CD4$^{–}$, Tim-4$^{–}$CD4$^{+}$, and Tim-4$^{+}$CD4$^{+}$ macrophage subsets at each time point. # P ≤ 0.01, § P ≤ 0.001, † P ≤ 0.0001. **(B)** Statistical comparisons were performed with a paired t test. **, P ≤ 0.01; ***, P ≤ 0.001; ****, P ≤ 0.0001.

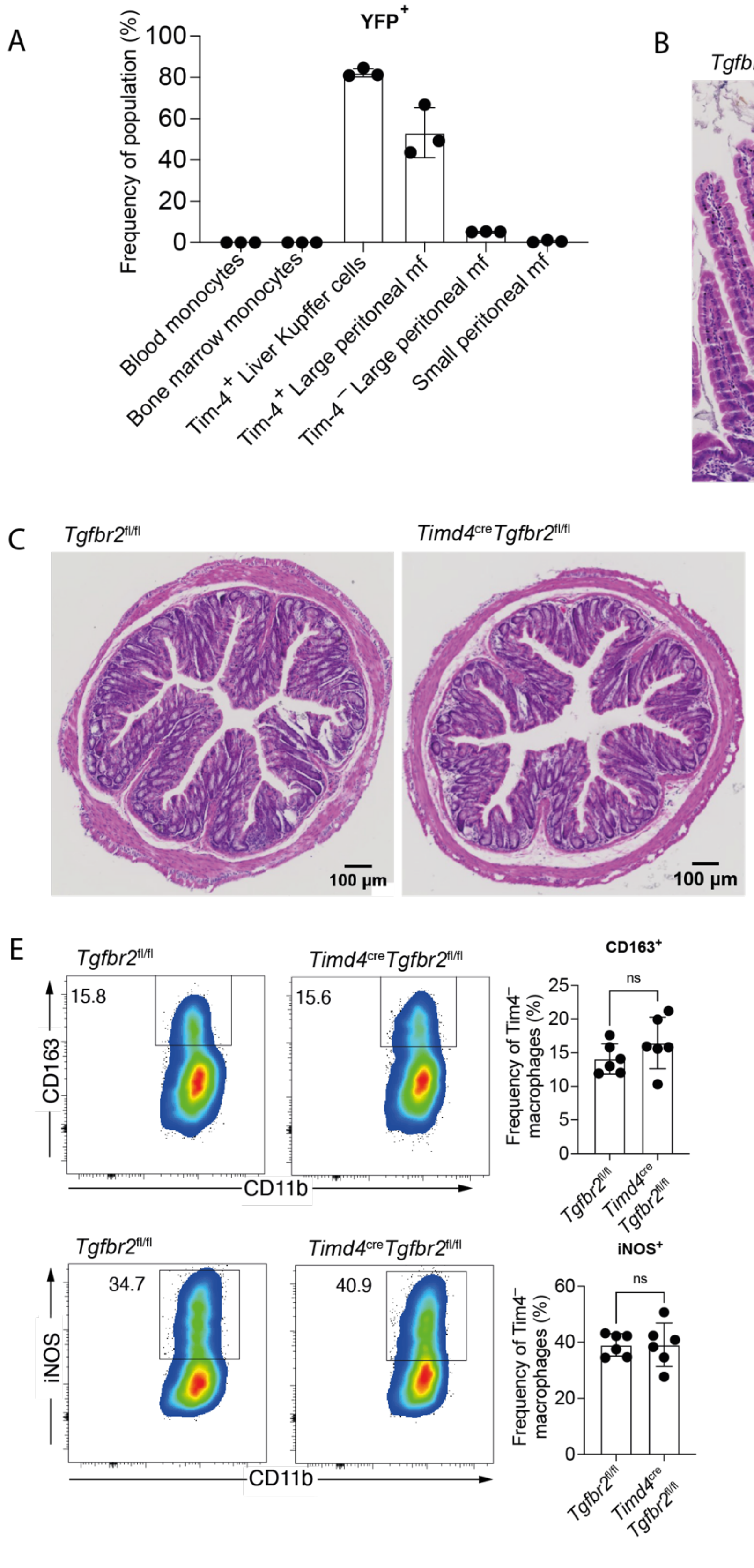


**Supplementary Figure 4. Characterisation of novel *Timd4*cre mouse line.**

**(A)** Frequency of Ly6C$^{hi}$ blood and bone marrow monocytes, Tim-4$^{+}$ liver Kupffer cells, Tim-4$^{+}$ large peritoneal macrophages, Tim-4$^{-}$ large peritoneal macrophages and small peritoneal macrophages expressing YFP. Data (n = 3) are from a single experiment. **(B)** Representative haematoxylin & eosin-stained small intestine from *Timd4*$^{cre}$*Tgfbr2*$^{fl/fl}$ mice and *Tgfbr2*$^{fl/fl}$ mice. **(C)** Representative haematoxylin & eosin-stained colon from *Timd4*$^{cre}$*Tgfbr2*$^{fl/fl}$ mice and *Tgfbr2*$^{fl/fl}$ mice. **(D)** Frequency of Tim-4$^{+}$CD163$^{-}$ and Tim-4$^{+}$CD163$^{+}$ small intestinal macrophages expressing iNOS in response to overnight LPS and IFNγ stimulation. **(E)** Upper left: representative flow cytometry plots showing the frequency of Tim-4$^{-}$ macrophages expressing CD163. Upper right: frequencies of Tim-4$^{-}$ macrophages expressing CD163. Lower left: representative flow cytometry plots showing the frequency of Tim-4$^{-}$ macrophages expressing iNOS. Lower right: frequencies of Tim-4$^{-}$ macrophages expressing iNOS. **(D, E)** Data (n = 5 per group) are pooled from 2 independent experiments. Error bars show mean ± SD. Statistical comparisons were performed with an unpaired t test with Welch's correction. *, $P \leq 0.05$.

| Cluster 1 genes |
|---|
| *1810011O10Rik, 2310057J18Rik, 2510009E07Rik, 2610008E11Rik, 2610035D17Rik, 2610203C22Rik, 2610507B11Rik, 4930539E08Rik, 5031425E22Rik, 5930403N24Rik, 9030617O03Rik, A4galt, AB124611, Abca1, Abca3, Abca9, Abcc5, Abcd2, Abi2, Ablim1, AC125444.1, AC152979.5, AC153955.2, Acsl1, Acss1, Acta2, Actg2, Adam15, Adamts10, Adap2, Adcy9, Adgre1, Adgre5, Adgrg6, Adk, Adrb2, Afap1l1, Aff3, Agmo, Agpat2, Agr2, Agtrap, Ahnak, AI427809, AI467606, Aicda, Akap12, Akap13, Aldh1a1, Aldh2, Aldh7a1, Alox5, Alox5ap, Ampd3, Ang4, Ankrd46, Anks1, Anln, Aoah, Ap1b1, Ap2a2, Apbb1, Apobec3, Apold1, App, Arap3, Arc, Arf2, Arhgap12, Arhgap22, Arhgap45, Arhgef10, Arhgef10l, Arhgef3, Arhgef6, Arid5a, Arl13b, Armc5, Arntl, Arrb1, Asap1, Asb10, Atf3, Atp2a3, Atp6v0a1, Atp6v0d2, AW549877, AY761184, B3galnt1, B3gnt7, B430306N03Rik, B4galt6, Bag2, Bag3, Batf, Bbc3, BC005537, BC035044, Bcl2l1, Bcl3, Bcl6, Bcl7a, Bend5, Bin1, Birc3, Blk, Blnk, Bmp2, Bmp2k, Bmpr1a, Bmyc, Btnl2, C1qtnf1, C1ra, C1rl, C1s1, C330027C09Rik, C3ar1, C4b, C5ar1, C5ar2, C7, Cables1, Calcrl, Camk1, Camkk2, Capn2, Capsl, Car13, Casp4, Cass4, Cav2, Cbr2, Ccdc126, Cchcr1, Ccl11, Ccl12, Ccl2, Ccl3, Ccl4, Ccl6, Ccl7, Ccl8, Ccl9, Ccnb1, Ccnd1, Ccnd3, Ccr6, Cd14, Cd163, Cd19, Cd209f, Cd209g, Cd300ld, Cd300ld2, Cd36, Cd37, Cd38, Cd55, Cd79a, Cd79b, Cd84, Cd93, Cdc42ep2, Cdca2, Cdca7, Cdk14, Cdk6, Cdkn1c, Cdkn2aip, Cdkn2c, Cdkn2d, Cdr2, Cebpd, Cenpa, Cenpe, Cep112, Cep68, Cep85, Ces1d, Cfh, Cfp, Cftr, Ch25h, Chd3os, Chka, Chp2, Chst12, Chst3, Chst7, Chsy1, Cited2, Ckap2, Ckb, Clec10a, Clec1a, Clec4a1, Clec4a3, Clmp, Cltc, Clu, Cmah, Cnksr3, Cnn3, Col14a1, Col1a1, Col1a2, Col3a1, Col6a1, Col6a2, Colca2, Colec12, Cp, Cpd, Cpe, Cpne8, Cps1, Cr2, Cracr2b, Creb5, Crim1, Crnde, Csad, Csf1, Csrp2, Cst3, Ctdsp2, Ctla2b, Ctnna1, Ctnnd1, Ctps2, Ctsd, Cx3cr1, Cxcl12, Cxcl13, Cxcr5, Cxxc5, Cyb5a, Cyfip1, Cyr61, Cyth4, D1Ertd622e, Dab2, Dag1, Dcaf12, Dclk2, Dclre1c, Dcn, Ddah2, Ddit3, Defa20, Defa21, Defa22, Defa24, Defa30, Defa32, Defa34, Defa5, Dek, Dennd2c, Des, Dgkd, Dgkh, Dhrs3, Diaph2, Disc1, Dmpk, Dnaja1, Dnajb1, Dnajb4, Dnajb5, Dnajc9, Dnmt1, Dock11, Dock8, Dpt, Dpysl2, Dpysl3, Dse, Dsn1, Dtl, Dtna, Dusp1, Dusp16, Dusp3, Dusp7, Dusp8, E2f7, Ebf1, Ednrb, Egr1, Egr2, Ehd4, Eif1a, Elk4, Emp3, Eng, Ephx1, Epn2, Epop, Eps15, Eps8, Esco2, Espl1, Esr1, Ets1, Ets2, Etv1, Etv3, Etv5, Exoc6b, F13a1, F630028O10Rik, F830016B08Rik, Fam102b, Fam111a, Fam118a, Fam129a, Fam135a, Fam160a2, Fam199x, Fam208a, Fam20c, Fam234a, Fam43a, Fas, Fbln1, Fbxo30, Fbxo4, Fbxo45, Fcer2a, Fcgr2b, Fcgr3, Fcgrt, Fcmr, Fcna, Fcrla, Fcrls, Fem1b, Fer, Fez2, Fgd6, Fggy, Fhl1, Fignl1, Filip1l, Fkbp1a, Fkbp1b, Fkbp4, Fkbp5, Fkbp9, Fli1, Flot2, Fmo2, Fnbp1l, Fndc7, Folr2, Foxp1, Foxp2, Foxred2, Fpr1, Frmd6, Fscn1, Fut10, G530011O06Rik, Gab1, Gab3, Gabpb1, Gadd45g, Gas6, Gas7, Gatm, Gatsl2, Gbp6, Gcc1, Gcsam, Gdf15, Gem, Gimap3, Gimap4, Gimap6, Glul, Gm13479, Gm13710, Gm14636, Gm15232, Gm15931, Gm16867, Gm20559, Gm21188, Gm26532, Gm26740, Gm29340, Gm30329, Gm34225, Gm37352, Gm44250, Gm44751, Gm4951, Gm5086, Gm5431, Gm7694, Gm8995, Gna15, Gpr165, Gpr34, Gpr84, Gprc5c, Gpt2, Gramd3, Grap, Grk6, Gtse1, Gypc, H2-Eb2, H2-K2, H2-Q5, H2-Q6, H2-Q7, H2afv, Hand1, Hbb-bs, Hbegf, Hdac10, Hdac5, Hdac9, Heca, Hivep2, Hmgn2, Hmox1, Hnmt, Homer2, Hpgd, Hpn, Hs3st1, Hs3st3a1, Hs6st1, Hsp90aa1, Hspa1a, Hspa1b, Hspb1, Hsph1, Icam1, Id2, Ier2, Ier3, Ier5, Ifi206, Ifi207, Ifi208, Igf1, Igfbp4, Ighd, Ighg1, Ighg3, Ighm, Ighv1-58, Ighv1-59, Ighv1-74, Ighv1-9, Ighv10-3, Ighv11-2, Ighv14-3, Ighv2-4, Ighv6-3, Ighv7-3, Ighv8-2, Igkv1-132, Igkv14-111, Igkv16-104, Igkv4-80, Igkv4-91, Igkv6-32, Iglc2, Iglc3, Iglv1, Iglv2, Igsf9, Iigp1, Ikzf3, Il16, Il1rl1, Il21r, Il22, Il2rb, Il4, Il6ra, Ildr2, Incenp, Ing2, Ints6l, Iqgap2, Irak3, Irf1, Irf2bp1, Irf2bpl, Irgm2, Irgq, Itga6, Itgb1, Itln1, Itm2b, Itpkc, Itsn1, Jag1, Jun, Kank2, Kbtbd7, Kcnip3, Kcnj9, Kctd12, Kctd21, Kdelc2, Kdr, Khnyn, Kif1b, Kif22, Kif23, Kif2c, Kif3a, Kitl, Klf10, Klf11, Klf2, Klf5, Klf6, Klhl5, Klrb1-ps1, Krt19, Krt8, Lamc1, Layn, Lbr, Lfng, Lgals1, Lgals2, Lifr, Lig1, Lilr4b, Lilrb4a, Lima1, Lmna, Lonrf3, Lpar1, Ltb, Ltc4s, Lum, Ly6d, Lyl1, Lyve1, Lyz2, Lzts2, Mad2l1, Maf, Maged1, Maml2, Man1a, Maoa, Map3k8, Map7d3, Mapk6, Marcks, Marcksl1, Mat2a, Mcc, Mcl1, Mcm2, Mcm3, Mcm6, Mctp1, Mef2c, Metap1d, Metrnl, Mgl2, Mgp, Mgst1, Mid1ip1, Miga1, Mir99ahg, Mmp9, Mob3c, Mpp1, Mrc1, Mrvi1, Ms4a1, Ms4a4d, Msh2, mt-Co1, mt-Nd1, mt-Nd2, mt-Nd4, Mt1, Mt2, Mtmr10, Mtmr12, Mtss1, Myh10, Myh11, Myl2, Myl9, Mylip, Mylk, Myo18a, Naa50, Nacc2, Nbl1, Ncald, Ncaph, Nck2, Ncl, Ndfip1, Necap2, Nectin4, Negr1, Neil3, Nes, Neurl1a, Nexn, Nfatc2, Nfic, Nfix, Nfkb2, Nfkbia, Nfkbib, Nfkbid, Nfkbiz, Nfxl1, Nhlrc3, Nhsl2, Nid2, Ninj1, Nkx2-3, Nme4, Npnt, Npy, Nr1d1, Nr1d2, Nr3c1, Nrp1, Nrp2, Nrros, Nt5c2, Nuf2, Nxpe5, Nxt2, Nynrin, Oas2, Optn, Osbpl11, Osm, Otc, Otub2, Otud1, Oxct1, P2rx1, P2rx7, P2ry12, P2ry6, P3h2, P4ha2, Pacsin2, Palld, Paqr9, Pbk, Pbx3, Pcgf6, Pcp4l1, Pcyox1, Pcyox1l, Pcyt1b, Pde2a, Pde4d, Pdlim1, Pdlim3, Pdp1, Pea15a, Peak1, Pepd, Per2, Pf4, Pgc, Phf11a, Phf11b, Phf19, Phlda1, Phlda3, Phyhd1, Pid1, Pigr, Pik3cg, Pkd2, Pkmyt1, Plcb3, Pld2, Plekhg2, Plekhg5, Plekhn1, Plk1, Plk4 ,Plpp2, Plppr4, Pls3, Plscr4, Pltp, Plvap, Plxna4, Plxnb2, Pml, Pmp22, Pogk, Pole, Polg, Pon3, Pou2af1,* |

| |
|---|
| *Pou2f2, Ppbp, Ppp1r10, Ppp1r9a, Prdx4, Prkar1b, Prkar2b, Prkcq, Prmt2, Pros1, Prps2, Prune2, Psd3, Psrc1, Pstpip1, Ptafr, Ptger4, Ptgr1, Ptk2, Ptov1, Ptpdc1, Ptprm, Ptpro, Qk, Rab11fip5, Rab12, Rab31, Rab6b, Rab7b, Rac2, Rac3, Rad21, Rad51b, Ralgps2, Ramp1, Rapgef4, Rapgef5, Raph1, Rasa4, Rasgrp2, Rasgrp3, Rasgrp4, Rassf4, Rbpj, Rcan1, Rcn1, Rcsd1, Reg3b, Reg3g, Rela, Relb, Rell1, Reps2, Retnla, Rfx2, Rgl1, Rgl3, Rgs13, Rgs3, Rhob, Rhobtb1, Rhoc, Ripor2, Rnase4, Rnf141, Rnf144b, Rnf145, Rnf150, Rnf185, Rock2, Rora, Rpa2, Rtn4, Rtp4, Ryk, S1pr1, Samd4, Samd9l, Sardh, Sbf2, Sbno2, Scarf1, Scd1, Scgn, Scn1b, Sdc3, Sdc4, Selenbp1, Selenop, Sell, Sema4a, Sema6b, Sept8, Serinc3, Serpina3f, Serpina3n, Serpinb6a, Serpinb8, Serpinf1, Serping1, Serpinh1, Sertad1, Sesn1, Sfmbt1, Sfrp1, Sft2d2, Sgcb, Sgce, Sh3bgrl2, Sh3bp5, Siah2, Siglec1, Siglech, Sirt1, Ski, Slamf9, Slc12a5, Slc12a7, Slc13a3, Slc14a1, Slc16a10, Slc20a1, Slc25a10, Slc25a24, Slc25a37, Slc35e4, Slc39a8, Slc41a2, Slc5a3, Slc7a13, Slc8b1, Slco2b1, Slfn10-ps, Slfn2, Slfn5, Slfn8, Slfn9, Smagp, Smc3, Smc4, Smo, Smpd1, Smpdl3b, Sms, Snap47, Snta1, Snx2, Snx6, Snx8, Socs3, Sorbs3, Sox4, Sparc, Spats2l, Sphk1, Spib, Spic, Spred1, Spry1, Sqstm1, Srpk3, Ssbp2, Ssh2, Sst, St3gal6, St6gal1, Stab1, Stap1, Stard3nl, Stard8, Stat3, Steap3, Steap4, Stmn1, Stom, Stxbp5, Sult1a1, Sun1, Susd1, Susd3, Swap70, Syne2, Synj2, Syt3, Tagap, Tagln2, Tanc2, Tank, Tbxas1, Tceal1, Tceal8, Tcf19, Tcf21, Tcf4, Tcof1, Tead2, Tef, Tgfbr2, Tgif2, Ticam1, Tifa, Timp2, Timp3, Tle1, Tln2, Tlr2, Tlr4, Tlr5, Tm6sf1, Tmcc1, Tmcc2, Tmem107, Tmem119, Tmem176a, Tmem176b, Tmem64, Tmem71, Tmem8, Tmem88, Tmpo, Tnf, Tnfaip8l2, Tnfrsf13c, Tnfsf12, Tnfsfm13, Tnip2, Tob1, Topors, Tox2, Tpbgl, Tpcn1, Tpi1, Tpm1, Tpm2, Tppp, Tpst1, Tpx2, Trem2, Trf, Trim24, Trim32, Trim47, Trim59, Trim8, Trove2, Trp53i11, Trps1, Trpv4, Tspan18, Tspan3, Tspan4, Tspan9, Ttk, Tuba1b, Tubb2a, Tubb6, Tubgcp5, Tuft1, Ube2f, Ubtd1, Uhrf1, Unc13b, Usp7, Vcam1, Vim, Vmp1, Vnn3, Vrk2, Wdhd1, Wls, Wsb1, Wtip, Wwp1, Wwtr1, Xylt2, Ypel5, Ywhaq, Zbtb10, Zbtb16, Zbtb2, Zbtb46, Zc3h12a, Zcchc24, Zeb1, Zfp367, Zfp36l1, Zfp422, Zfp512, Zfp703, Zfp704, Zmym6* |
| Cluster 2 genes |
| *1600014C10Rik, 1700108F19Rik, 2010300C02Rik, 2210408F21Rik, 2900026A02Rik, 6530402F18Rik, 9530059O14Rik, A4gnt, A830008E24Rik, Aacs, Abca7, Abcb1a, Abcg1, Abcg3, Abi3, Abr, Abt1, AC160562.1, Acly, Acp5, Acvrl1, Acy1, Adam19, Adam23, Adamtsl4, Adap1, Adgre4, Adgrg5, Adgrl3, Adpgk, Adprh, Aga, Agap1, Agpat5, Ahcyl1, Ahr, Aif1, Aifm1, Ak2, Akr1a1, Aldh1b1, Aldoa, Aldoc, Alg1, Alg2, Amacr, Amz1, Ankrd66, Antxr1, Apobr, Apoc2, Apol10b, Apol7b, Apol7c, Arf3, Arf4, Arhgap27, Arl11, Arl4c, Arl5a, Arl6ip1, Arpc3, Arpin, Asb2, Atg101, Atg4c, Atp13a1, Atp13a3, Atp1a3, Atp2a2, Atp5d, Atp6v0e, Atp6v1b2, Atp6v1e1, Atp6v1f, Atp6v1g1, Atrnl1, AU020206, AW112010, B230217C12Rik, B3galt5, B3gnt8, B4galt4, Bbs5, BC031181, Bcap29, Bckdha, Bcl2a1d, Bcl2l14, Bhlhe40, Bin2, Bpnt1, Brox, Bscl2, Btg2, Bvht, Bzw2, C130050O18Rik, C130089K02Rik, C2cd2l, Calm1, Capn10, Car2, Card11, Casp3, Casp6, Casp7, Cbl, Cbr3, Ccdc102a, Ccdc158, Ccdc166, Ccdc86, Ccdc88b, Ccdc97, Ccl24, Ccr1, Ccrl2, Ccz1, Cd164, Cd1d1, Cd200r1, Cd209e, Cd244, Cd274, Cd300a, Cd300c2, Cd300e, Cd300lf, Cd6, Cd80, Cd82, Cd9, Cdc42se2, Cdipt, Cebpzos, Cep152, Cep83, Cfb, Chd7, Chmp5, Chpf, Chuk, Cib1, Ciita, Cisd2, Cish, Clec5a, Clec7a, Clip2, Cln8, Cmtm8, Coa5, Colgalt1, Coq10b, Cox5a, Cox8a, Creg1, Crem, Crtc3, Csf2ra, Csf2rb, Csf2rb2, Csnk1e, CT025556.1, Ctage5, Ctnnbl1, Ctsh, Ctsl, Ctss, Ctsz, Cwc25, Cxcl10, Cxcl16, Cxcl9, Cyb5r3, Cyfip2, Cyld, Cyp4f16, Cyp4f18, Cyp51, Cysltr2, Cytip, D16Ertd472e, D8Ertd738e, Dad1, Dbi, Dbnl, Dck, Ddhd1, Derl1, Dfna5, Dgat1, Dgat2, Dgke, Dhcr24, Dhcr7, Dhrs11, Dip2c, Dlst, Dnajb6, Dnajc16, Dnase1l3, Dock10, Dock5, Dpp4, Dtx4, Dym, Dync1li1, Dyrk4, Ece1, Efnb1, Eif2ak1, Eif2b2, Eif4g3, Elf4, Ell2, Emd, Entpd6, Epb41, Ephx2, Eprs, Ero1lb, Evl, Exoc3l4, F11r, F3, Fabp1, Fabp6, Fads2, Fads3, Fam105a, Fam117a, Fam160b2, Fam189a2, Fam234b, Fam26f, Fam46c, Fbl, Fbxl17, Fbxo32, Fbxo6, Fdps, Fgl2, Fgr, Flnb, Fndc5, Frrs1, Fth1, Fuca1, Fuca2, Fundc1, Fyn, Gad1-ps, Galm, Galnt12, Galnt6, Galnt7, Ganc, Gatsl3, Gbp2b, Gbp3, Gbp4, Gbp5, Gbp7, Gbp8, Gcnt2, Gde1, Gdf3, Gdi2, Gfpt1, Ggt5, Ggta1, Gk, Gkn3, Gla, Glrx, Gm14005, Gm15448, Gm15922, Gm19434, Gm20056, Gm2a, Gm32633, Gm33819, Gm37168, Gm37199, Gm37347, Gm37531, Gm38248, Gm45716, Gm5150, Gm6377, Gm8221, Gmppb, Gngt2, Gnl2, Gnptab, Golim4, Got1, Gpd1l, Gpr108, Gpr141, Gpr171, Gpr55, Gpr65, Gramd4, Grhl1, Grk3, Gsdmd, Gsr, Gsto1, Gstp1, Gtf2h2, H1f0, H2-T23, Hap1, Havcr2, Hcar2, Hck, Heatr1, Hic1, Hist1h1c, Hk2, Hlx, Hmgcl, Hmgcr, Hmgcs1, Hnrnpll, Hoxb4, Hoxb5, Hs3st3b1, Hsd17b12, Hsd17b4, Hsd17b7, Hsd3b7, Htatip2, Hvcn1, I830077J02Rik, Idh1, Idnk, Ifi44, Ifnar1, Igha, Ighj1, Ighv1-12, Ighv1-18, Ighv1-19, Ighv1-26, Ighv1-38, Ighv1-4, Ighv1-50, Ighv1-52, Ighv1-54, Ighv1-55, Ighv1-64, Ighv1-72, Ighv1-76, Ighv1-78, Ighv1-80, Ighv1-81, Ighv1-82, Ighv2-9-1, Ighv3-1, Ighv3-6, Ighv4-1, Ighv5-12, Ighv5-17, Ighv5-9, Ighv5-9-1, Ighv6-6, Ighv9-2, Igkc, Igkv1-110, Igkv1-117, Igkv1-135, Igkv1-88, Igkv10-94, Igkv12-41, Igkv12-44, Igkv12-46, Igkv13-85, Igkv17-121, Igkv17-127, Igkv19-93, Igkv2-109,* |

| |
|---|
| *Igkv3-10, Igkv3-12, Igkv3-7, Igkv4-50, Igkv4-57-1, Igkv4-59, Igkv4-63, Igkv4-72, Igkv5-37, Igkv5-39, Igkv5-43, Igkv5-45, Igkv5-48, Igkv6-13, Igkv6-14, Igkv6-15, Igkv6-17, Igkv6-20, Igkv6-23, Igkv6-25, Igkv8-18, Igkv8-21, Igkv8-24, Igkv8-27, Igkv8-30, Igkv9-120, Igsf6, Igsf8, Il12b, Il12rb2, Il13ra1, Il18bp, Il1b, Il1r2, Il1rl2, Il6st, Impa1, Impdh2, Inafm2, Insig1, Insl6, Ipo13, Irak2, Irf6, Irs2, Isg15, Isoc1, Itga1, Itga4, Itgal, Itgax, Itgb2, Itgb5, Itm2c, Jaml, Jchain, Jmy, Kazn, Kcnj10, Kcnk6, Kctd6, Khk, Kif9, Klhl18, Klra17, Klra2, Klrb1b, Kmo, Kpna4, Kynu, Lbh, Lcmt2, Ldha, Ldlr, Ldoc1l, Leprot, Lgals3, Lgals9, Lgmn, Lipa, Lipe, Lmtk2, Lpcat2, Lrch1, Lrp10, Lrrc4c, Lrrfip1, Lst1, Ltbr, M6pr, Mafb, Malt1, Maml3, Man2a1, Man2b1, Mansc1, Map4k3, Mapk13, Mapk8, Mapkapk3, march7, Mcfd2, Med7, Mefv, Met, Mfsd12, Mfsd13a, Mgat4a, Mgat5, Mical1, Midn, Mlkl, Mmd, Mmp14, Mocos, Mpeg1, Mpp6, Mpzl2, Mpzl3, Mrm1, Ms4a4a, Ms4a6b, Ms4a6c, Ms4a6d, Msmo1, Msrb1, Mtfr1, Mtpn, Muc6, Mxd1, Mycl, Nabp1, Naga, Nagk, Naip2, Nampt, Narf, Ncapg2, Nceh1, Ncf2, Ncf4, Ncoa7, Ndel1, Ndst1, Ndufa9, Nedd4l, Nedd9, Neurl3, Nfatc1, Nfil3, Nfkbie, Nif3l1, Nipal3, Nlrx1, Nod2, Nop9, Notch1, Npc2, Nr1h3, Nr4a3, Nsdhl, Ntn4, Ntpcr, Nuak2, Nucb2, Nudt18, Nup98, Nus1,Nxn, Nxpe4, Ocstamp, Olfml3, Ostm1, Ovca2, P2ry2, P4hb, Palm, Panx1, Pard6a, Pbxip1, Pcyt2, Pdcd1lg2, Pdcl3, Pde1b, Pde4b, Pdia4, Pdxk, Pecam1, Per1, Pex10, Pgf, Pgs1, Pianp, Pik3cb, Pik3r5, Pilra, Pilrb1, Pilrb2, Pim3, Pip5k1c, Pira2, Pkib, Pkm, Pla2g16, Pla2g4a, Pla2g7, Plaur, Plbd1, Plcb2, Plcl1, Plk3, Plpp5, Pls1, Plxdc1, Plxdc2, Pmepa1, Pmvk, Polr2g, Pparg, Ppfia4, Ppm1g, Ppm1h, Ppp2cb, Ppt1, Ppt2, Pram1, Prdm1, Prdx1, Prdx5, Preb, Prelid3b, Prosc, Prr5l, Prss30, Psap, Psd4, Psma1, Psmc1, Psmd14, Psme1, Psme2, Ptgs2, Ptk2b, Ptms, Ptp4a1, Ptpn22, Ptpn7, Ptprc, Ptprs, Pvrig, Pxdc1, Rab11fip4, Rab19, Rab32, Rab4b, Rap2a, Rapgef1, Rasal3, Rasgrp1, Rbck1, Relt, Rfc2, Rgs1, Rgs12, Rgs2, Rheb, Rhog, Ric1, Rin3, Ripk3, Rnasek, Rnd3, Rnf115, Rnf149, Rnf43, Rnft1, Rogdi, Rsad1, Rsad2, Rspo1, Rundc3b, Runx2, Runx3, Rusc1, S100a1, Samd8, Sap18, Sc5d, Scap, Scarb1, Scarb2, Scd2, Scel, Scimp, Sdf2l1, Sdhaf2, Sdhd, Sec13, Selenof, Selenok, Selenom, Sema4b, Sept11, Sgk1, Sh2d1b1, Sh3bp1, Sharpin, Sidt2, Siglecf, Sik1, Sirpb1b, Sla, Slc15a3, Slc16a6, Slc25a13, Slc25a20, Slc25a3, Slc25a33, Slc26a2, Slc2a1, Slc30a1, Slc35c2, Slc35d2, Slc35e1, Slc37a3, Slc39a6, Slc3a2, Slc44a2, Slc44a5, Slc7a11, Slc7a7, Slc9a3r1, Slco3a1, Slco4a1, Smad6, Smad7, Smim3, Smox, Snhg15, Snx10, Snx18, Snx20, Soat1, Socs2, Socs6, Sorl1, Spg21, Sphk2, Spi1, Spint1, Spon1, Spsb1, Spty2d1, Sqle, Src, Srsf10, Ss18l1, Ssfa2, Ssr4, Stard4, Stat1, Stbd1, Stk17b, Stk38l, Ston2, Stra6l, Stx2, Stxbp2, Sulf2, Sumf1, Sys1, Taldo1, Tapbp, Tapbpl, Tbc1d1, Tbc1d9, Tcp11l2, Tctn3, Tep1, Tff2, Tfip11, Tgfb1, Tgfbi, Tgfbr1, Tgm2, Thbs1, Themis2, Tifab, Tinf2, Tiparp, Tjp2, Tlr12, Tlr13, Tm2d2, Tm4sf19, Tmc6, Tmeff1, Tmem131, Tmem14c, Tmem150b, Tmem156, Tmem206, Tmem268, Tmem50b, Tmem51, Tmem97, Tmx1, Tmx3, Tnfaip2, Tnfrsf11a, Tnfrsf1a, Tnfrsf1b, Tnfrsf21, Tomm34, Tor1a, Tor1b, Tpst2, Treml4, Trim6, Trit1, Trmt10c, Trmt61b, Tsc22d1, Tsg101, Tspan13, Tspan33, Tuba4a, Tyrobp, Ubald1, Ube2j2, Ubl3, Ubxn8, Ucp2, Uevld, Unc45a, Unc93a, Usmg5, Usp12, Usp36, Usp6nl, Vamp4, Vhl, Vill, Vipas39, Vipr1, Vopp1, Vps37b, Wipf1, Wnt4, Wsb2, Yipf3, Zbtb7b, Zc3h12d, Zfp366, Zfp386, Zfp622, Zfp667, Zfp958, Zfyve28, Zfyve9, Zscan20, Zyg11b* |
| Cluster 3 genes |
| *2900052N01Rik, 4632427E13Rik, 4732496C06Rik, 4833411C07Rik, Abca5, Abcc3, Abcd4, Abhd12, Abhd16a, Abl1, AC165247.1, Acp2, Acpp, Acsf2, Actr3b, Adam22, Adam28, Adam33, Adamdec1, Adcy4, Adcy7, Adgb, Adgra3, Adgrl2, Adh6a, Adipor1, Adora3, Adrb1*<br>*Agpat3, Ahrr, Akr1b10, Aldh1l1, Aldh3b1, Aldob, Angptl4, Ankrd16, Ap3m2, Ap5s1, Ap5z1, Apoc1, Appl2, Arel1, Arhgap10, Arhgap19, Arhgap4, Arhgef18, Arhgef5, Arid5b, Arl4d, Armc8, Arrdc3, Art2a-ps, Art2b, Aspa, Atp13a2, Atp2b2, Atp8a1, AW011738, Axl, B4galnt4, Bank1, Basp1, Baz2a, BC037034, Bco2, Birc6, Blvrb, Brat1, Btnl7-ps, C1qa, C1qc, C2, C6, Cadm1, Camk2d, Camk2n1, Cant1, Capn3, Casc4, Casp12, Catip, Ccnl2, Cd200r4, Cd209b, Cd22, Cd4, Cd63, Cd72, Cd81, Cdc42bpa, Cdk11b, Cdk16, Cdkn1a, Cebpa, Cebpg, Cfap74, Chic1, Chpf2, Chst14, Ckmt1, Clasp2, Clcn7, Clec1b, Clec4n, Clk4, Clock, Clstn1, Cmklr1, Cmtm4, Cndp2, Cpeb4, Cped1, Cpq, Creg2, Csf1r, Ctc1, Ctns, Ctsf, Cttn, Cxcl1, Cxcl14, Cyb561d1, Cyb5r1, Cyp27a1, Cystm1, Cyth1, Cyth3, D7Ertd128e, Dapk3, Dennd2a, Dgki, Dkk3, Dlc1, Dmxl2, Dnaaf3, Dnah2, Dnajb2, Dnajc28, Dnase2a, Dock4, Dok1, Dok3, Dst, Dtnbp1, Dtx3, Dusp6, Ebi3, Echdc2, Ecm1, Edil3, Eef2k, Efemp2, Ehf, Elavl4, Engase, Enpp1, Enpp2, Enpp4, Enpp5, Eogt, Epb41l2, Epb41l3, Epg5, Epm2aip1, Epor, Ermap, Esam, Eva1b, Eya4, F2r, F830208F22Rik, Fabp2, Fads1, Fam13a, Fam213a, Fam46a, Fap, Fblim1, Fbp2, Fbxo21, Fcgr4, Fchsd2, Fcrl1, Firre, Fmn1, Fmnl2, Fmo5, Fnip2, Fos, Frmd4a, Fsd2, Fstl4, Fxyd2, Fzd8, Gaa, Galnt3, Gas1, Gbgt1, Gdap10, Gdi1, Gdpd1, Gkap1, Gm12958, Gm13391, Gm13431, Gm13994, Gm14221, Gm15880, Gm15964, Gm26520, Gm26917, Gm26947, Gm28042, Gm43682, Gm44860, Gna11, Gna12, Gns, Gpd1, Gpm6b, Gpnmb, Gpr137b, Gpr137b-ps, Gpr157,* |

| |
|---|
| *Gpr176, Gpr31b, Gpx3, Grina, Gstm2, Gstm3, H2-M2, H6pd, Hacd3, Hes1, Hip1, Hist1h2bc, Hist3h2a, Hjurp, Hk3, Hmox2, Hpdl, Hpgds, Hrh1, Hs1bp3, Hs2st1, Hunk, Icosl, Idua, Ifi27, Ifit3, Ifit3b, Igfbp3, Ighv1-15, Ighv1-39, Ighv10-1, Ighv5-6, Igkv1-133, Igkv10-96, Igkv14-126, Igkv15-103, Igkv3-4, Igkv3-5, Igkv4-54, Igkv4-55, Igkv4-56, Igkv4-68, Igkv4-78, Il10, Il12rb1, Il27, Impact, Inpp4a, Inpp4b, Inpp5j, Insr, Irf2bp2, Irf8, Ispd, Itgav, Itpkb, Jade2, Jtb, Kansl3, Kcng2, Kcnj16, Kcnk13, Kcp, Kctd7, Keap1, Kifc3, Klhl13, Klhl41, Klhl6, Klhl9, Krt20, Lag3, Lamp1, Lao1, Laptm4a, Laptm4b, Leng8, Lgals3bp, Lgals4, Lgals8, Lgr4, Lilra5, Lix1, Lmo3, Lncpint, Loxl3, Lpin1, Lrp6, Lrrc57, Lyplal1, Mab21l3, Mag, Malat1, Man1c1, Mapre3, March1, Marveld2, Max, Mb21d2, Mbd4, Meis1, Meis3, Mertk, Mfge8, Miip, Mlh3, Mmp10, Mmp13, Mmp2, Morc3, Mospd2, Mr1, Mras, Mroh2a, Ms4a14, Ms4a7, Msr1, mt-Cytb, mt-Rnr1, Mthfr, Mtus1, Mxi1, Myo1a, Myo1e, Myo7a, Myo9a, Naglu, Nbr1, Nckap5, Nckap5l, Neat1, Necap1, Neil1, Neo1, Nfat5, Nisch, Nlrc4, Nlrp1c-ps, Nod1, Npepps, Npl, Nr2f6, Nudt16, Oasl1, Ocln, Ogt, Olfr111, Olfr1330, Olfr561, Ophn1, Oplah, Osbpl10, Osgin1, P2rx4, P2rx6, P2ry13, P4ha1, Pald1, Pde8b, Pdgfb, Pdgfc, Peg13, Pgap1, Phactr1, Phf23, Phospho2, Pigz, Pik3c2a, Pik3r1, Pik3r3, Pim1, Pitpnc1, Pkp4, Pla2g15, Pla2g2d, Pla2g4b, Plagl2, Plcd1, Pld3, Plekha8, Plekhb2, Plekhm2, Plin2, Plk2, Plod1, Pnisr, Pnpla7, Pnrc2, Pomk, Postn, Ppcdc, Ppfibp2, Ppp1r21, Ppp2r2b, Prag1, Prkab1, Prpf38b, Prr5, Pstpip2, Ptgs1, Ptpn13, Ptpn14, Pyroxd2, Rab33b, Rab34, Rab3il1, Rassf1, Rbm7, Rbp2, Rcbtb2, Renbp, Rgl2, Rgmb, Rgs10, Rhbdf1, Rhbdf2, Rhobtb3, Rhoh, Rims3, Rin2, Rnasel, Rnf103, Rnf180, Rnf186, Rnf215, Rps4l, Rspry1, Rufy3, Rusc2, Rxra, Sash1, Sat1, Scamp5, Scarf2, Scly, Scrn3, Sec14l1, Selenon, Sema4c, Sema6d, Serpina3g, Sertad3, Sgpl1, Sgsh, Sh3bp2, Sh3bp4, Sh3d19, Sh3d21, Sipa1l1, Sirt7, Slc11a1, Slc12a2, Slc1a3, Slc22a17, Slc26a11, Slc28a2, Slc29a1, Slc31a2, Slc35a5, Slc37a2, Slc38a9, Slc40a1, Slc43a2, Slc46a1, Slc4a8, Slc7a4, Slc7a8, Slc8a1, Slc9a3r2, Slc9a9, Slf2, Slit3, Smc1b, Smim1, Snx11, Snx24, Snx29, Soga1, Spats2, Specc1l, Speg, Spg20, Spire1, Spock1, St14, St5, St6galnac3, Stab2, Stag3, Stap2, Stard13, Sult1b1, Sult1d1, Svbp, Tanc1, Taz, Tbc1d10a, Tbc1d12, Tbc1d23, Tbcc, Tcaf1, Tcf7l2, Tcirg1, Tcn2, Tesk2, Tet2, Tex264, Tfec, Thap12, Thnsl2, Thrb, Tigd2, Timd4, Tk2, Tmem104, Tmem106a, Tmem140, Tmem2, Tmem221, Tmem26, Tmem37, Tmem55b, Tmem82, Tmem86a, Tmem87b, Tmem9b, Tmprss5, Toe1, Tor4a, Tpcn2, Tpp1, Trafd1, Triap1, Trim25, Trp53inp2, Trpm2, Tsc22d3, Tsku, Tspan15, Tspan8, Tssc4, Ttyh1, Ttyh2, Txlnb, Txnip, Uaca, Uba7, Ulk2, Unc5b, Use1, Usp21, Vps41, Vsir, Wdr20, Wdr81, Wdr91, Wfdc17, Whamm, Xdh, Xlr, Ythdc1, Zc3h7a, Zdhhc14, Zdhhc24, Zfp110, Zfp263, Zfp281, Zfp472, Zfp623, Zfp641, Zfp69, Zfp691, Zfp707, Zfp715, Zfp773, Zfp820, Zfp839, Zfp84, Zfp992, Zfyve27, Znfx1, Zscan26* |
| Cluster 4 genes |
| *1700025G04Rik, 2210010C04Rik, 2310022A10Rik, 4833407H14Rik, 5430437J10Rik, A530064D06Rik, A630033H20Rik, Abcc4, Abce1, Abracl, AC133083.2, AC163354.1, AC238811.2, Acap1, Acod1, Acot11, Acot7, Actg1, Actn1, Actr2, Actr3, Adam8, Add3, Adora2a, Adora2b, Adss, Adssl1, Afap1, Afdn, Agpat4, AI504432, AI506816, AI839979, Akr7a5, Alcam, Aldh1a2, Alpk2, Alyref, Amot, Amy2b, Anp32b, Anp32e, Anpep, Anxa1, Anxa2, Ap1s2, Apba1, Aqp9, Areg, Arhgap26, Arhgdib, Arhgef37, Arl2bp, Arl5c, Arl6ip5, Arpc1b, Arpc2, Arpc4, Arpc5, Arsb, Asf1b, Ass1, Atox1, Atp10a, Atp11b, Atp1a1, Atp5c1, Atp5e, Atp5g1, Atp5g3, Atp8b4, Atxn10, Aurka, Aurkb, Avpi1, Azin1, B3gnt5, B4galnt1, B4galt5, Bach1, Bambi, Banf1, Bcl11a, Bcl2a1a, Bcl2l11, Bend4, Birc5, Bri3bp, Btf3, Btg1, Btla, Bub1, Bub1b, C1qbp, C3, Calm3, Capg, Capzb, Cbfb, Ccdc12, Ccl17, Ccl22, Ccna2, Ccnb2, Ccnf, Ccr2, Ccr7, Cct5, Cd101, Cd209a, Cd24a, Cd2ap, Cd300lb, Cd300lg, Cd44, Cd52, Cd69, Cd7, Cdc14a, Cdca3, Cdca7l, Cdh1, Cdk2ap2, Cdv3, Cela1, Cenpf, Cenpl, Cenpm, Cep55, Cers6, Cfl1, Chil3, Chn2, Chp1, Ckap2l, Cks1b, Cks2, Clec12a, Clec4a4, Clec4b1, Clec4b2, Clec4e, Clic1, Clic4, Clps, Cmas, Cnbp, Cnn2, Copg2, Copz1, Coq2, Coro1a, Coro2a, Cotl1, Cox7a2l, Cpm, Crip1, Cs, Csgalnact2, Csrp1, Cstb, CT010467.1, Ctnnd2, Ctrl, Cxcl3, Cxcr4, Cyb561a3, Cyb5r4, Dapk1, Dcstamp, Ddb1, Ddr1, Ddx21, Dennd3, Dennd4a, Dhx40, Diaph3, Dmkn, Dna2, Dpep2, Dpy19l1, Dusp2, Dusp5, Dut, Dynll1, E2f2, Ear2, Eef1a1, Eef1g, Eef2, Egr3, Eif3b, Eif3e, Eif3f, Eif3h, Eif3i, Eif3k, Eif3l, Eif3m, Eif4b, Eif5a, Elovl5, Emb, Emilin2, Eml2, Emp1, Endod1, Eno1, Enoph1, Epcam, Erlin1, Ero1l, Errfi1, Esyt1, Etf1, Exosc5, Ezr, F10, Fabp5, Fam107b, Fam110a, Fam49b, Fam69a, Fam96a, Far1, Fasn, Fau, Fbxo34, Fcho1, Fem1c, Ffar2, Fgfr1, Fgfr1op, Flna, Flrt3, Flt3, Fmnl1, Fn1, Fosl2, Frat2, Furin, Fxyd5, G3bp1, G6pdx, Gabarapl2, Gapdh, Gapt, Gbp2, Gcat, Gch1, Gcsh, Gda, Glipr1, Glipr2, Glud1, Gm12159, Gm13373, Gm1673, Gm1966, Gm43814, Gm44135, Gm5424, Gm8113, Gnai2, Gnas, Gng10, Gosr2, Got2, Gp2, Gpr132, Gpr35, Gpr4, Gprc5a, Gpx1, Grasp, Gsn, Gtf2b, H2-DMb2, H2-Oa, H2-Ob, H2afx, H2afy, H2afz, H3f3a, Haao, Hdc, Helz2, Hepacam2, Hexim1, Hif1a, Hilpda, Hmgb2, Hnrnpa1, Hopx, Hp, Hpcal1, Hpse, Hspbp1, Htr7, Id3, Idh3a, Ifi205, Ifi27l2a, Ifi30, Ifitm1, Ifitm3, Ifitm6, Ifngr1, Ighv1-14, Ighv1-21-1, Ighv2-2, Ighv7-1, Igkv13-84, Igkv14-100, Igkv3-2, Igkv4-61, Igkv9-123, Il17ra,* |

| |
|---|
| *Il18rap, Il1a, Il1rn, Il22ra2, Il23a, Il6, Il7r, Impa2, Inhba, Ipcef1, Ipo5, Ipo7, Irf4, Isg20, Itga5, Itgae, Itgam, Itgb7, Itpr1, Jak2, Jarid2, Jdp2, Jpt1, Kcna3, Kcnab2, Kcnd1, Kctd17, Kif11, Kit, Klk1, Klk8, Klrd1, Klri1, Klri2, Klrk1, Kmt5a, Krt80, Lactb, Ldhb, Ldlrad3, Lilra6, Limd1, Lin54, Lmnb1, Lmo4, Lpcat4, Lpl, Lpp, Lrrc32, Lrrc59, Lrrk2, Lsp1, Lsr, Lta4h, Ltb4r1, Ly6a, Ly6c2, Ly6i, Ly75, Lyz1, Maff, Map2k3, Map3k14, Map4k1, Map4k4, Mapk14, Mbnl3, Mbp, Mcemp1, Mcm4, Mcm5, Mcm7, Mcub, Mdh2, Me2, Med13l, Megf9, Mfsd6, Mfsd7b, Mif, Mir155hg, Mir17hg, Mki67, Mlec, Mmp12, Mmp25, Mob3a, Mob3b, Mogs, Mospd4, Mreg, Mrpl33, Ms4a4b, Ms4a4c, Ms4a8a, Msn, Mthfd1l, Mthfd2, Myc, Mycbp2, Myh9, Myl6, Myo1f, Myo1g, Naaa, Nab1, Naca, Napsa, Nav1, Nbeal2, Ncf1, Ndc80, Ndrg1, Ndufa4, Nedd4, Nedd8, Net1, Nfkb1, Nhp2, Nlrp3, Noc2l, Noct, Nop58, Npm1, Nr4a2, Nrg1, Nucks1, Nup210, Nupr1, Nusap1, Oas3, Odc1, Ola1, Olfm1, Olr1, Osbpl3, Osgin2, P2ry10, Pa2g4, Pabpc1, Padi2, Paics, Pak1, Parp1, Parvg, Pcdh1, Pdk3, Pfkp, Pfn1, Pglyrp1, Plac8, Plcg2, Plcxd2, Plekha5, Plet1, Plscr1, Plxnc1, Plxnd1, Pnlip, Pnliprp1, Pnliprp2, Polr1a, Polr2e, Ppa2, Ppie, Ppm1m, Ppp1ca, Ppp1r14a, Ppp1r1a, Pptc7, Pqlc3, Prdx6, Prep, Prkag2, Prkar2a, Prkd3, Prkx, Prmt5, Procr, Prss2, Psat1, Psma4, Psma7, Ptger3, Ptma, Ptpa, Pvr, Pxylp1, Pygl, Qpct, Rab11fip1, Rab27a, Rab5c, Rabgef1, Racgap1, Rack1, Rai14, Ramp3, Ran, Ranbp1, Rbbp7, Rbpms, Rcc1, Rcc2, Reg1, Rel, Rgcc, Rgs18, Rhov, Rmi2, Rnase1, Rnase6, Rnf217, Rnh1, Rpl10, Rpl11, Rpl13a, Rpl14, Rpl15, Rpl17, Rpl18, Rpl18a, Rpl19, Rpl21, Rpl22, Rpl23, Rpl23a, Rpl26, Rpl27, Rpl29, Rpl3, Rpl30, Rpl32, Rpl34, Rpl35, Rpl36, Rpl36a, Rpl37, Rpl37a, Rpl39, Rpl4, Rpl41, Rpl6, Rpl7, Rpl7a, Rpl8, Rpl9, Rplp0, Rplp1, Rplp2, Rps10, Rps11, Rps13, Rps14, Rps15, Rps15a, Rps16, Rps18, Rps19, Rps2, Rps20, Rps23, Rps24, Rps25, Rps26, Rps27, Rps27a, Rps27l, Rps3, Rps3a1, Rps4x, Rps5, Rps6, Rps6ka2, Rps7, Rps8, Rps9, Rpsa, Rrad, Rras2, Rrm1, Rrm2, Rrp12, Rrp1b, S100a10, S100a11, S100a6, Samsn1, Sdc1, Sec24d, Sec61b, Seh1l, Selplg, Sem1, Sema4d, Sema7a, Sept6, Sept9, Serpinb1a, Serpinb6b, Serpinb9, Serpini2, Set, Sf3b6, Sgk3, Sgms1, Sh2d3c, Sh3bgrl3, Shcbp1, Siglecg, Sirpb1a, Sirpb1c, Skint3, Slamf7, Slamf8, Slc16a1, Slc16a3, Slc1a5, Slc2a6, Slc36a3os, Slc38a1, Slc38a2, Slc41a1, Slc46a3, Slc52a3, Slc9a7, Slfn1, Smpdl3a, Snrpe, Snrpf, Sod2, Sort1, Sowahc, Sp100, Spc24, Spc25, Spint2, Spns3, Spp1, Srebf2, Srgap3, Srgn, Srsf7, Srxn1, Ssrp1, St3gal1, St3gal4, St3gal5, St8sia6, Stil, Stk10, Stk24, Stt3b, Stx11, Stx3, Sub1, Sumo1, Svip, Syngr2, Taf13, Taf4b, Tbc1d4, Tbx21, Tctex1d2, Tes, Tfdp1, Thbd, Tiam1, Timeless, Tkt, Tle3, Tlr11, Tmed9, Tmem123, Tmem38b, Tmsb10, Tmx4, Tnfaip8l1, Tnfrsf12a, Tnfrsf13b, Tnfrsf18, Tnfsf8, Tnfsf9, Tnip3, Tnni2, Top2a, Tpm4, Tppp3, Traf1, Traf4, Trem1, Trem3, Trerf1, Trib1, Try4, Tspan2, Tspo, Tspoap1, Ttc39c, Ttc7, Tuba1a, Tuba1c, Tubb5, Twf2, U2af1, Ube2c, Uck2, Unc119b, Utp18, Vars, Vasp, Vcl, Vdr, Vegfa, Vrk1, Wdfy4, Wdr1, Wnt11, Xcr1, Xylt1, Ybx3, Zbtb18, Zc3h12c, Zc3hav1, Zdhhc13, Zg16, Zmiz2, Zswim4, Zyx* |

**Supplementary Table 1**
